# Concomitant processing of choice and outcome in frontal corticostriatal ensembles correlates with performance of rats

**DOI:** 10.1101/2020.05.01.071852

**Authors:** Takashi Handa, Rie Harukuni, Tomoki Fukai

## Abstract

The frontal cortex-basal ganglia network plays a pivotal role in adaptive goal-directed behaviors. Medial frontal cortex (MFC) encodes information about choices and outcomes into sequential activation of neural population, or neural trajectory. While MFC projects to the dorsal striatum (DS), whether DS also displays temporally coordinated activity remains unknown. We studied this question by simultaneously recording neural ensembles in the MFC and DS of rodents performing an outcome-based alternative choice task. We found that the two regions exhibited highly parallel evolution of neural trajectories, transforming choice information into outcome-related information. When the two trajectories were highly correlated, spike synchrony was task-dependently modulated in some MFC-DS neuron pairs. Our results suggest that neural trajectories concomitantly process decision-relevant information in MFC and DS with increased spike synchrony between these regions.

## Introduction

Deciding actions based on the outcome of past actions is crucial for the survival of animals. The frontal cortex-basal ganglia network is engaged in various forms of decision making involved in adaptive goal-directed behaviors [1-4]. These studies have shown how choice- and/or outcome-related information is processed in the frontal cortex and the basal ganglia. However, how the cortico-basal ganglia network encodes behavioral information at the neural ensemble level remains largely unknown. How the related brain regions communicate such information during outcome-based action selection is yet to be clarified [5]. In this study, we explore whether and how task-related neural activity is coordinated between the medial frontal cortex (MFC) and dorsal striatum (DS), which is a major input structure of the basal ganglia [6-8], of rats performing an outcome-based alternative choice task.

The MFC is engaged in encoding reward and error signals [9,10] and hence is considered to play a role in monitoring positive and negative outcomes from an action. In particular, the rostral agranular medial cortex or the secondary motor cortex of rodent has been implicated in sensory-cued [11-13] and outcome-based action selection [14,15]. Along the frontal cortico-basal ganglia axis, excitatory MFC outputs project monosynaptically to DS [16,17], which also receives glutamatergic inputs from some thalamic nuclei [7,18] and are modulated by midbrain dopaminergic inputs conveying outcome information [19] and reward prediction error [20]. Accordingly, DS is thought to associate a specific action with the resultant outcome [21-24]. Consistent with this, lesions in MFC [14,15] and DS [25,26] impair outcome-based choice behaviors.

Evidence from MFC, or more specifically posterior parietal cortex (PPC) and orbitofrontal cortex (OFC), shows that sequential activation of cortical neuron ensembles, which is termed neural trajectory, underlies decision making [12,27,28]. We explore how this trajectory information is conveyed to striatal neurons during decision making. If the role of the striatum is selection of cortical inputs for action generation, as conventionally thought, such selection may occur sparsely in time, making continuously evolving trajectories less relevant to striatal functions. We therefore clarify to what extent cortical and striatal neuron ensembles are coherently activated during decision making. We hypothesize that neural ensembles in the MFC and DS utilize synchronized spikes to communicate with one another during adaptive behavior. Such spikes are ubiquitous in cortical circuits [29,30] and were suggested to enhance information transmission and association between directly or indirectly interconnected brain regions [31-32].

We trained head-restrained rats to lick either left or right spouts for earning reward (positive outcome) and simultaneously recorded multi-neuron activity from the MFC and DS of these rats. Because a rewarded spout was changed intermittently, the rats had to switch their choice responses to maximize reward, exhibiting behavioral responses similar to a typical win-stay lose-away behavior.

Unexpectedly, we found that neural trajectories emerge and evolve highly coherently in both MFC and DS during the entire period of the alternative choice task. In some MFC-DS neuron pairs, the number of coincident spikes was modulated by task events when the trajectory evolution displayed an enhanced coherence. Our results suggest that the adaptive control of outcome-based decision making relies on the coevolution of neural trajectories in MFC and DS and that spike synchrony plays a role in this temporal coordination.

## Results

### Adaptive outcome-based behavior under head-restrained condition

Twenty head-restrained rats performed an outcome-based two choice task, in which the rats were required to lick one of two spouts (Left and Right) and in each block of trials one spout delivered reward, with a high reward acquisition probability (>75%, see Methods) (Fig.1a). The choice-reward contingency was systematically reversed without any sensory feedback after the cumulative number of rewarded trials reached 10 in each block. The rats figured out the reversal of choice-reward contingency through the monitoring of no-reward events, time-out, and reward acquisition within subsequent several trials (Fig. 1b). Rats tended to select the same spout as that chosen in the last rewarded trial, but they switched the choice following one to several unrewarded trials (Fig. 1b, c). This choice pattern is close to the so-called ‘win-stay’ and ‘lose-shift’ which is the optimal strategy to maximize reward in the present task. To examine if the choice pattern (repeating post rewarded trial and switching post no-reward trial) was biased by past outcome (reward or no-reward) or by past choice position (Left and Right), we estimated the effect of past outcomes and past choices by a logistic regression analysis (Methods). The last outcome provided the greatest effect on the current choice type (repeat or switch) regardless of past choice position (t-test with Bonferroni correction, p = 3.73×10^−15^). In contrast, the current choice position was the most affected by the last choice position regardless of outcome (t-test with Bonferroni correction, p = 5.17×10^−6^) (Fig. 1d).

**Figure 1.**
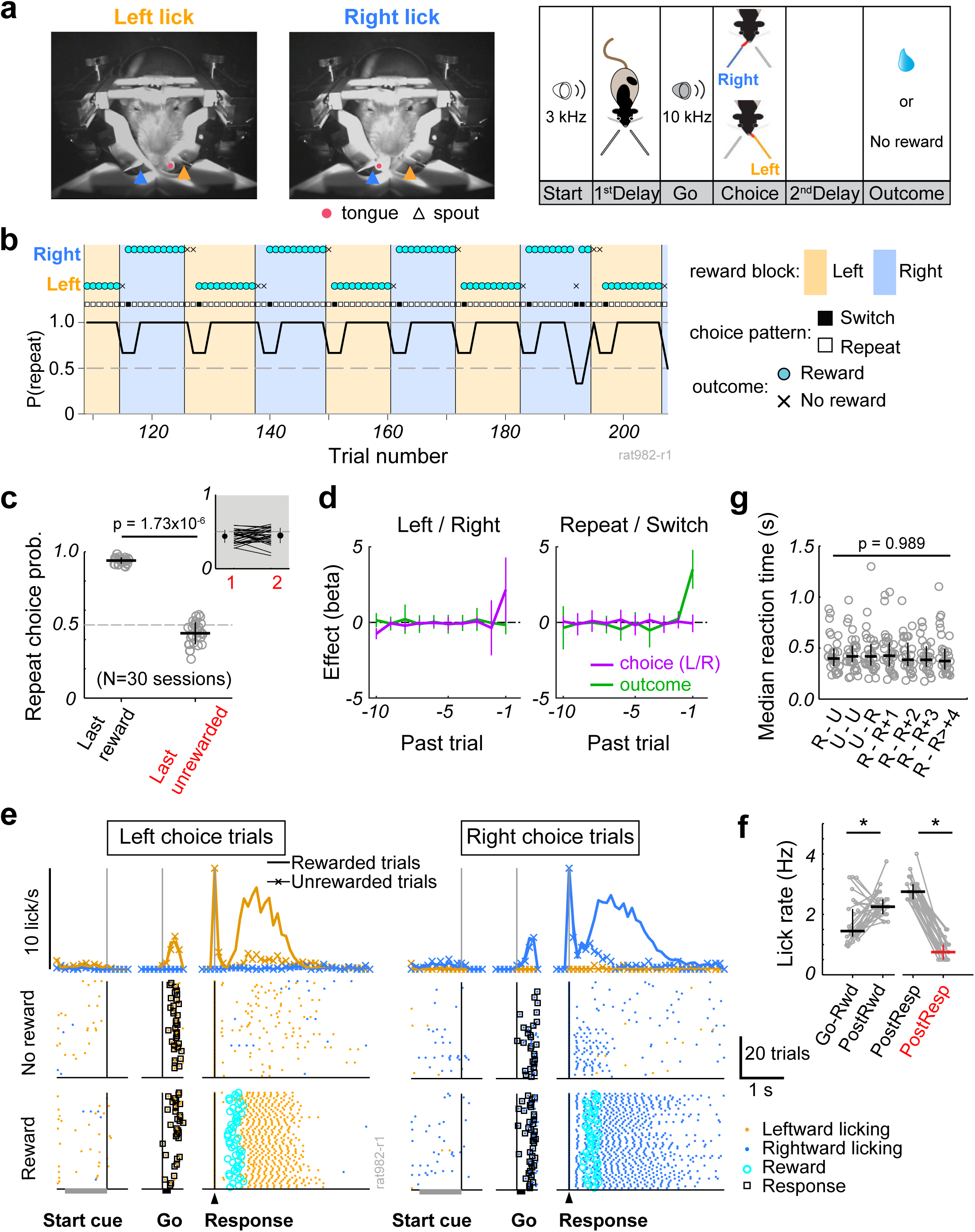
Adaptive choice patterns in response to change in choice-reward contingency. (a) *Left and middle*: Snapshots of a head-restrained rat making lick responses toward Left (orange arrowhead) and Right (blue arrowhead) spouts. A circle indicates the tongue position. *Right*: A schematic illustration of an outcome-based two alternative choice task. Individual trial started with ‘Start’ tone (3 kHz) presentation. Head-fixed rats were required to wait for ‘Go’ tone (10 kHz) without a lick during ‘1st Delay’ period, and then to make a choice response by licking either one of spouts within 5 s after ‘Go’ tone onset. Reward was given after ‘2nd Delay’ of various durations (0.3-0.7 s) if the chosen position corresponded to current reward location. Otherwise no reward, no sensory feedback, and timeout of 5 s were given. (b) Representative flexible choice behavior by a rat across 9 block-reversals of reward position. Vertical lines and colored areas denote the block-reversal and reward position in each block, respectively. Circle and cross indicate the outcome, reward and no-reward respectively, after choice response (Left or Right). Square shows choice pattern (Switch or Repeat). (c) Probability of repeating choice after rewarded and unrewarded outcomes (30 sessions, 20 rats). Circle indicates individual sessions. Horizontal lines and error bars represent mean and SD, respectively. The statistical significance was confirmed by Wilcoxon signed-rank test. Inset: Probability of repeating choice following one or two consecutive unrewarded trials. (d) Effects of past choice (purple) and outcome (green) on (*left*) current choice position and (*right*) current choice type (repeat and switch). The ordinate shows averaged coefficients of a logistic regression model (N = 30 sessions). Error bars show SD. (e) Licking patterns during rewarded and unrewarded trials in the session shown in b. Rastergrams denote licking times and locations (Left: orange, Right: blue). Histograms show averaged Left and Right licking rates in rewarded (solid line) and unrewarded (line with cross) trials. (f) Median licking rate in an epoch between Go tone onset and reward delivery (Go-Rwd), during outcome period of 4-s after reward delivery (PostRwd), and for 4-s after response in rewarded (PostResp in black) and unrewarded (PostResp in red) trials. Asterisks indicate statistical significance (Wilcoxon signed-rank test, p <0.05). (g) Comparison of reaction times among different choice patterns. Circles denote median reaction times in individual trials. Horizontal and vertical lines indicate median and 1st-3rd quartiles across trials, respectively. Trials were classified by last and current outcomes at each session (e.g., U–R denotes unrewarded last and rewarded current choices, and R+1, R+2, … denote the number of continuous rewarded outcomes).

Trial-by-trial licking behavior was strongly influenced by outcome events. Rats licked either one of the two spouts immediately after the appearance of Go tone, and then showed fast and rhythmic licking at around 6 Hz just after reward delivery (Fig. 1e). In contrast, rats showed sparse licking at around 1 Hz during outcome period in unrewarded trials. The lick rate of population data was significantly different before and after reward delivery in rewarded trials (Fig. 1f, Wilcoxon signed-rank test, p = 0.00295) and between rewarded and unrewarded outcome periods (Wilcoxon signed-rank test, p = 1.50×10^−6^). In behavioral studies on primates and rodents, prolonged response times, called post-error slowing, are observed after no reward outcome in sensory-cued choice tasks [33,34]. At 13 sessions, we observed significantly differential reaction times among different choice patterns classified by recent outcome history (Kruskal-Wallis test, p <0.05). Although the reaction times were indeed longer after no or first reward outcome than after consecutive reward outcomes in some sessions, we did not find any tendency toward post-error slowing at the population level (p = 0.989, Kruskal-Wallis test, n = 30 sessions) (Fig. 1g).

### Simultaneous multi-neuron recordings from MFC and DS

To test whether and how neuronal population activities of the MFC-DS circuit are collectively coordinated during the outcome-based action selection, we recorded multi-neuron activity in both MFC and DS using two silicon probes during task performance. In addition, we simultaneously performed juxtacellular recordings of Neurobiotin-loaded neurons near the silicon probe for the MFC recording (Fig. 2a). We confirmed the projection from the recording site in the MFC to that in the DS by using a retrograde tracer Fluoro-Gold (FG) injected into the DS (Methods). FG-labelled corticostriatal neurons were primarily observed in certain restricted regions of the MFC (Fig. 2b and c), particularly in the rostral agranular medial cortex as previously reported [6,7]. Observing the track of the silicon probe insertion, Neurobiotin-loaded neurons (in some cases), FG-labelled neurons in layers 3 and 5 of MFC (Fig. 2b-c and Fig. S1) and the tracks of another probe for the DS recording at or near FG injection site in DS (Fig. 2b), we confirmed that multi-neuron activity was presumably recorded from a subset of MFC-DS circuit.

**Figure 2.**
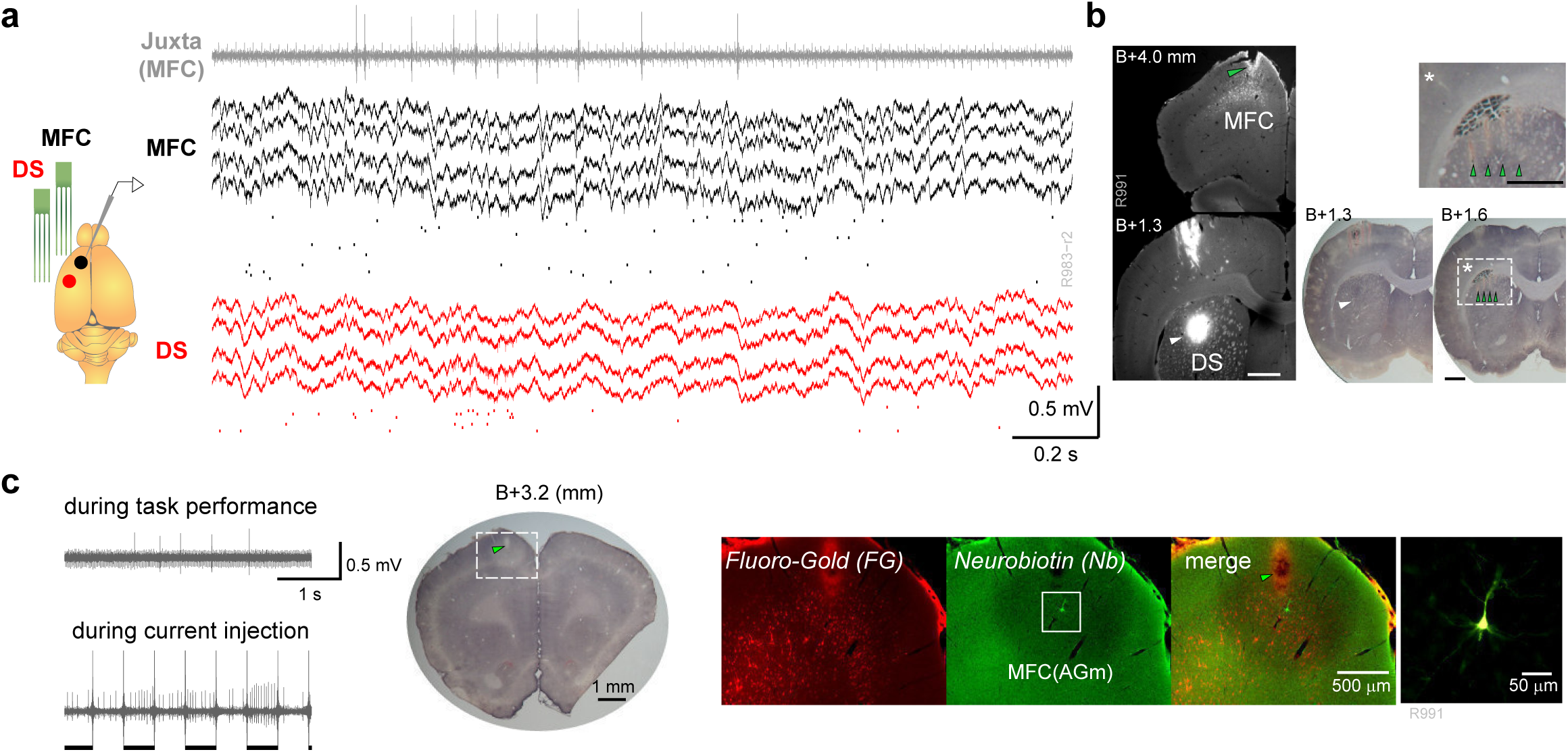
Simultaneous recording from MFC and DS. (a) Left: A schematic illustration of recordings. One silicon probe (black) and a glass pipette electrode (gray) were inserted into MFC and another silicon probe into DS (red). Right: Multiunit activity recorded with tetrodes in MFC and DS and juxtacellularly recorded activity (gray). Spikes of isolated units are depicted below multiunit signals. (b) Left: Fluorescent images show corticostriatal projection neurons labelled with Fluoro-Gold (FG) in MFC (top) and the injection site (white arrowhead) of FG in DS (bottom). Right: Nissl stained brain sections specified by the AP coordinates from Bregma. One shown at bottom left is identical to the lower fluorescent image and one at bottom right indicates silicon probe tracks (scars, green arrowhead) in DS. The area surrounded by a dash-lined box was magnified (top). Scale bars show 1 mm. (c) Left: Juxtacellularly recorded activity of a neuron during task performance (top) and current injection (bottom). Horizontal bars indicate epochs of periodic current injection whereas long vertical lines are artifacts by the on/off cycles of injection. Middle: A Nissl stained brain section including probe track (scar, green arrowhead) in MFC (agranular medial cortex). Right: Fluorescent images reveal FG labelled corticostriatal neurons and a Neurobiotin (Nb)-labeled neuron in MFC. The rightmost image enlarges the inside of the box area.

**Figure S1.**
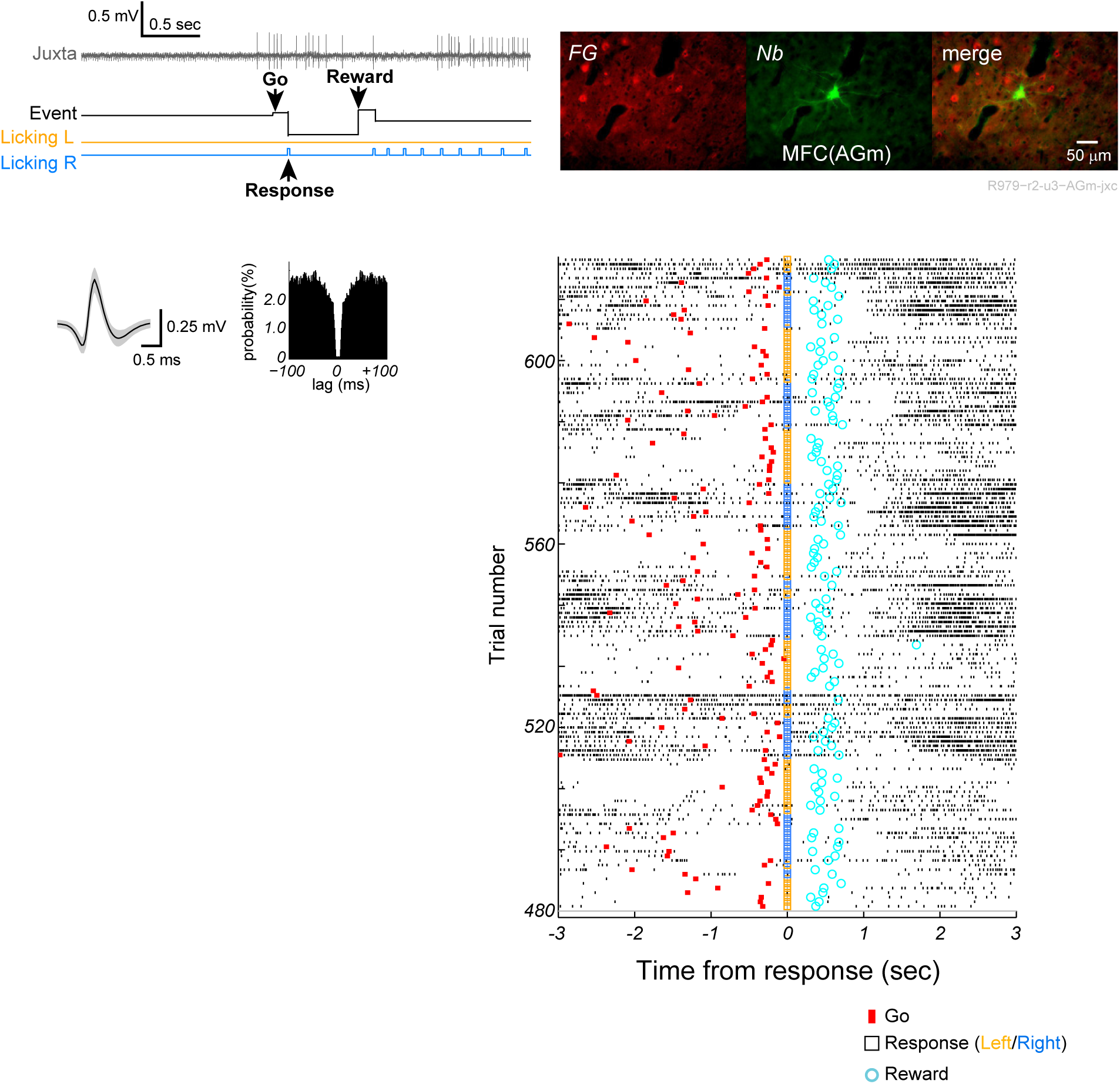
Juxtacellular recording during task performance and labelling the recorded neuron. Top left: Juxtacellular activity (gray) during task performance. Black, orange, and blue lines indicate event signal, left licking signal, and right licking signal, respectively. Top right: Post hoc visualization of the recorded neuron and corticostriatal projection neurons. Bottom left: Auto-correlogram and spike waveform of the identified neuron. Bottom right: Firing patterns of the labelled neuron aligned to response execution.

### Single MFC and DS cells encode both positive and negative outcomes

We analyzed event-related firing patterns of well-isolated single neurons recorded simultaneously from MFC (n = 468, mean ±SD = 39 ±18 cells per session) and DS (n = 489, mean ±SD = 40 ±14 cells per session) in 10 rats. The recordings were performed in the left hemisphere while these rats were performing the learned task through 12 recording sessions (Methods). The firing rates of both MFC and DS cells were correlated with choices (Left vs. Right) and outcomes (Reward vs. No-reward) (two-way ANOVA, p <0.05, Methods).

**Figure S2.**
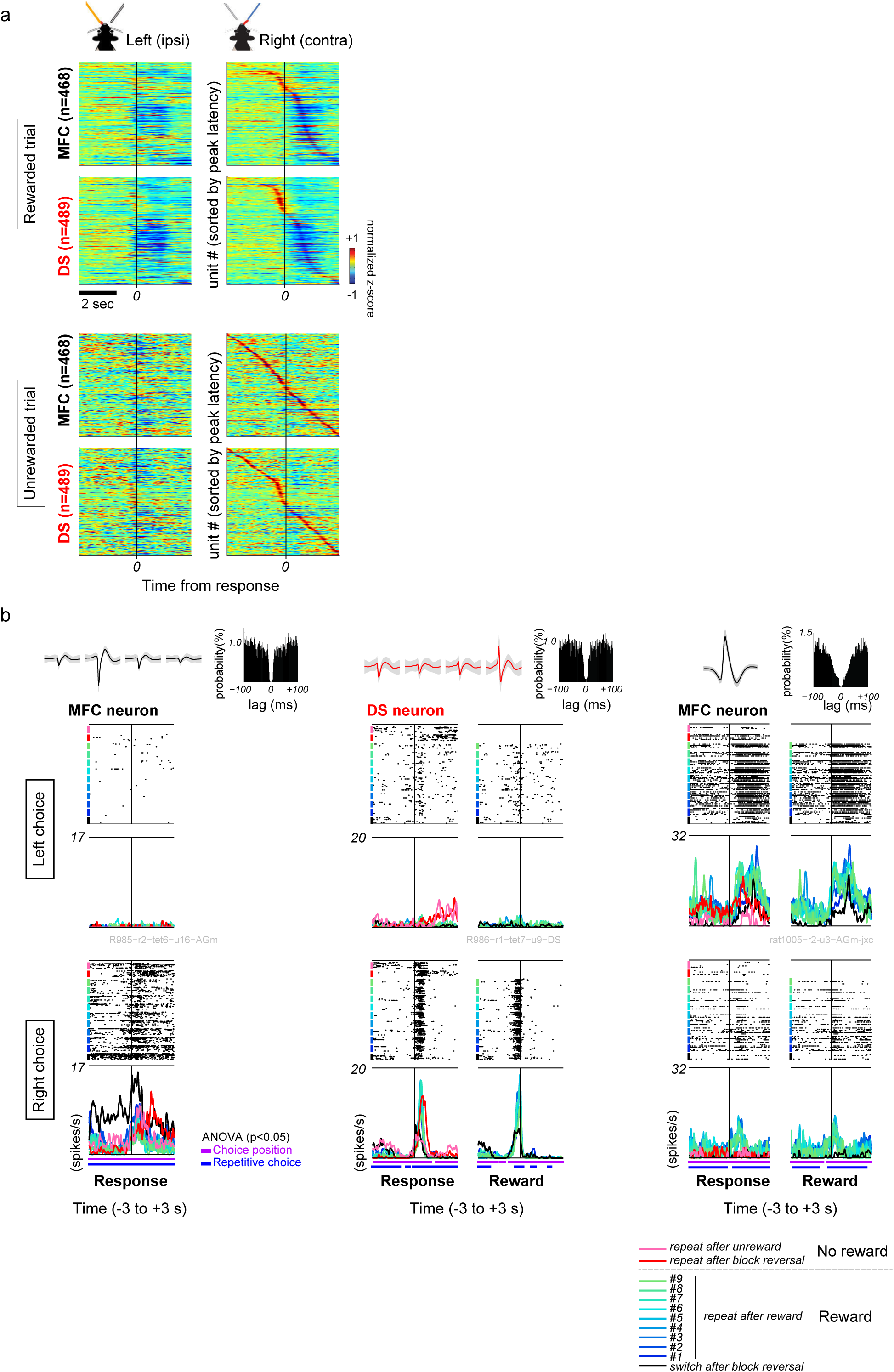
Complex feature of firing patterns in MFC and DS cells. (a) The same collective PETH as in Fig. 3e is presented in the order of units sorted by the latencies at maximum absolute amplitudes in rewarded (top) and unrewarded (bottom) right choice trials. (b) Three neurons changed firing rates between (left and middle) switch/repeat choices or (right) repetitive rewarded/unrewarded trials. Top row shows autocorrelations of spike time and spike waveforms recorded with tetrode (left and middle) and glass pipette (right). Trials are classified by the number of repetitive choices after rewarded and unrewarded outcomes. Rastergrams and PETHs are presented according to this classification. The PETHs denote trial-averaged firing rate. Horizontal bars under PETHs indicate times at which firing rate was significantly different between choice positions (purple) or among the number of repetitive choice trials (blue) (two-way ANOVA, p <0.05).

Many outcome-modulated neurons decreased firing rate during the period of positive outcome but not during that of negative outcome (Fig. 3a, b). However, some neurons increased firing rate most strongly for negative outcomes (Fig. 3c), and others raised firing rate for positive outcomes (Fig. 3d). We then calculated peri-event time histograms (PETHs), which revealed similar sequential dynamics of MFC and DS cells (Fig. 3e and Fig. S2a). The PETHs also indicated a prominent decrease in firing rate in the majority of MFC and DS cells after reward delivery. The fractions of choice- and outcome-modulated neurons evolved similarly in MFC and DS, but these fractions differed significantly between MFC and DS in large portions of task period (Fig. 3f). We quantified the degree and preference of selectivity across different sessions and neurons by performing receiver operating characteristic (ROC) analysis of neurons’ firing rates and calculating the area under ROC curve (AROC) (Methods). In both MFC and DS, choice-selective neurons exhibited a bias towards right (i.e., contralateral) choice responses whereas outcome-selective neurons towards unrewarded outcomes (Fig. 3f). At the group level, selectivity indices were significantly larger in MFC than in DS for both choice- and outcome-selective neurons in large portions of task period (Fig. 3g, Mann-Whitney U test, p <0.01). A small number of MFC and DS cells dynamically changed their firing patterns according to the histories of outcome and choice (Supplementary information and Fig. S2b).

**Figure 3.**
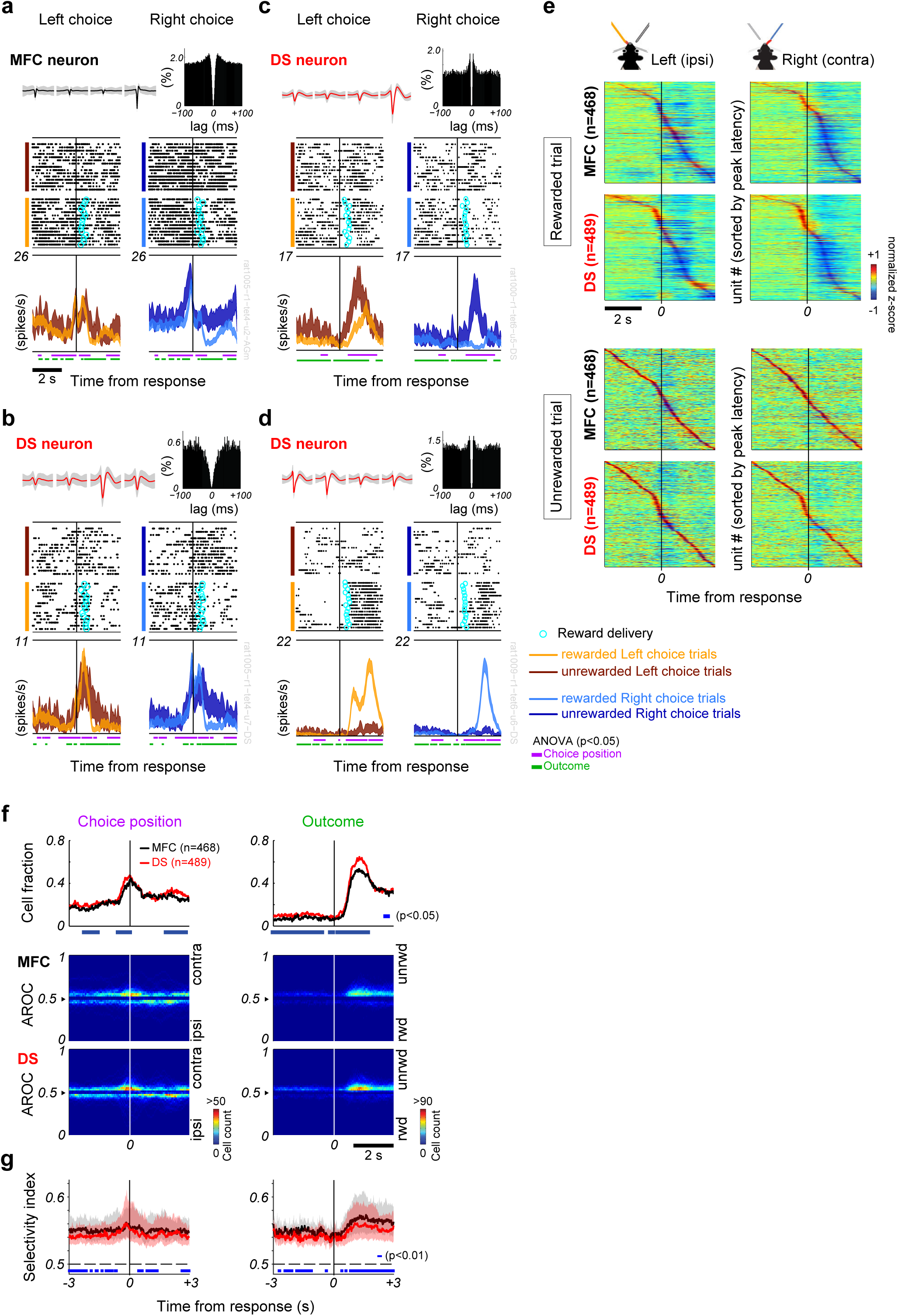
Choice and outcome coding in single neurons of MFC and DS. Below, neuronal activities are aligned to choice response (vertical line). (a-d) Representative firing patterns modulated by choices and outcomes on MFC and DS single neurons. Simultaneously recorded Right-choice selective responses of (a) MFC and (b) DS cells. These cells decreased firing rate immediately after reward delivery (cyan circle). Another DS cell increased firing rate during post-outcome periods in (c) unrewarded and (d) rewarded trials. Top row shows spike waveforms (mean ± SD) in four channels of a tetrode and auto-correlogram. Peri-event time histogram (PETH) shows mean ± SEM of firing rate (bin = 200-ms, sliding 20 ms), and horizontal bars indicate the times at which firing rate was significantly different (two-way ANOVA, p <0.05) with respect to choice (purple) or outcome (green) factor. (e) Normalized PETHs of MFC (n = 468) and DS (n = 489) neurons are shown in rewarded and unrewarded trials. In each condition, units are sorted by the latency of maximum absolute peaks. (f) *Top*: Proportion of MFC (black) and DS (red) cells showing (*left*) choice or (*right*) outcome-selective activity (two-way ANOVA, p <0.05). Blue bars represent the times at which the cell proportions were significantly different between MFC and DS (χ2 test, p <0.05). *Middle and bottom*: The number of choice- and outcome-selective neurons are shown together with AROC values revealing preference between contralateral/ipsilateral choices or unrewarded/rewarded outcomes, respectively. (g) Comparison of selectivity index for choice (*left*) and outcome (*right*) between MFC (black) and DS (red). Thick solid lines and shaded area represent median and 1st/3rd quartiles, respectively. Blue bars represent the times at which the selectivity indices were significantly different between MFC and DS (Mann-Whitney U test, p <0.01).

Thus, choice- and outcome-modulated neurons show similar dynamical evolution in MFC and DS, but their fractions and degrees of selectivity were significantly different between the two brain regions. These results suggest parallelism between MFC and DS in the neural population coding of task-relevant information in this outcome-based choice task.

### Functional correlation of neural trajectories between MFC and DS

The above similarity in neural dynamics raises a question about the way neural population activities are coordinated in MFC and DS to encode information about decision making. To clarify this, we explored how the neural trajectories encoding Left and Right decisions were separated in each session (12 sessions in total) by using Fisher’s linear discriminant (Methods). This method yields a hyperplane that optimally separates datasets belonging to two categories, such as Left/Right choices or positive/negative outcomes, in the high dimensional space of the instantaneous population rate vectors [12,13,35]. We calculated discriminability index *d’*, which is the distance between the categorized rate vectors measured orthogonally to such a hyperplane. To keep a sufficient number of cells (n ≧ 25 in both regions per session, Supplementary information), below we omitted three data sets (R986-r1, R1000-r1, R1004-r1 in Fig. S3a) and only used 9 datasets (R982-r1, R983-r1, R983-r2, R985-r1, R985-r2, R991-r1, R1005-r1, R1009-r1, R1012-r1).

**Figure S3.**
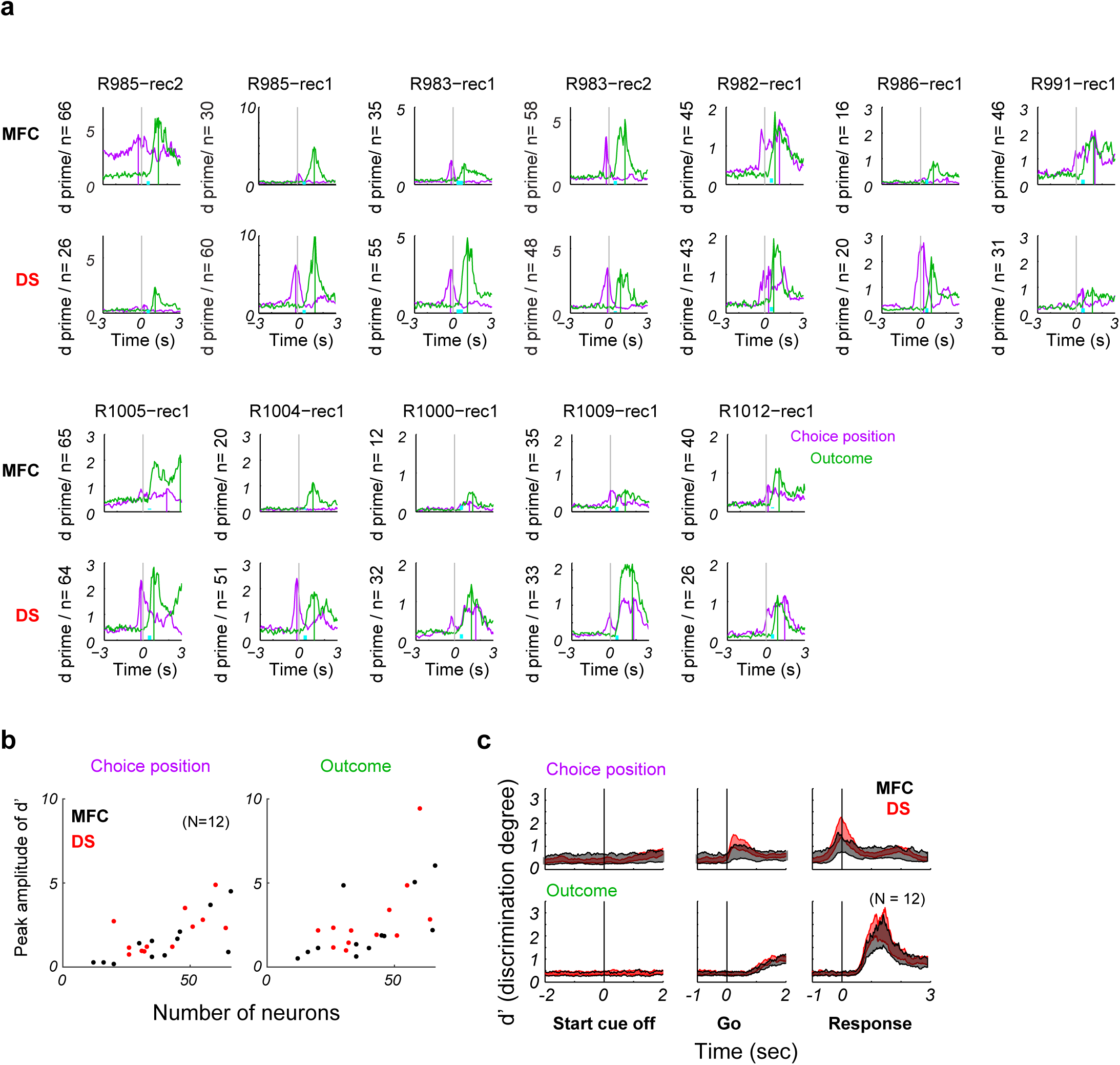
Degree of discrimination by population activity at individual sessions. (a) Time evolution of discrimination degree (d’) on choice (purple) and outcome (green) axes around choice responses. Vertical line indicates the peak time of d’ and cyan bar the interval of reward delivery. (b) Correlations between peak d’ values and the number of neurons in population data. (c) Group data of d’ on choice and outcome axes in MFC (black) and DS (red). Data were aligned at start cue end, go cue end, and response execution.

**Figure S4.**
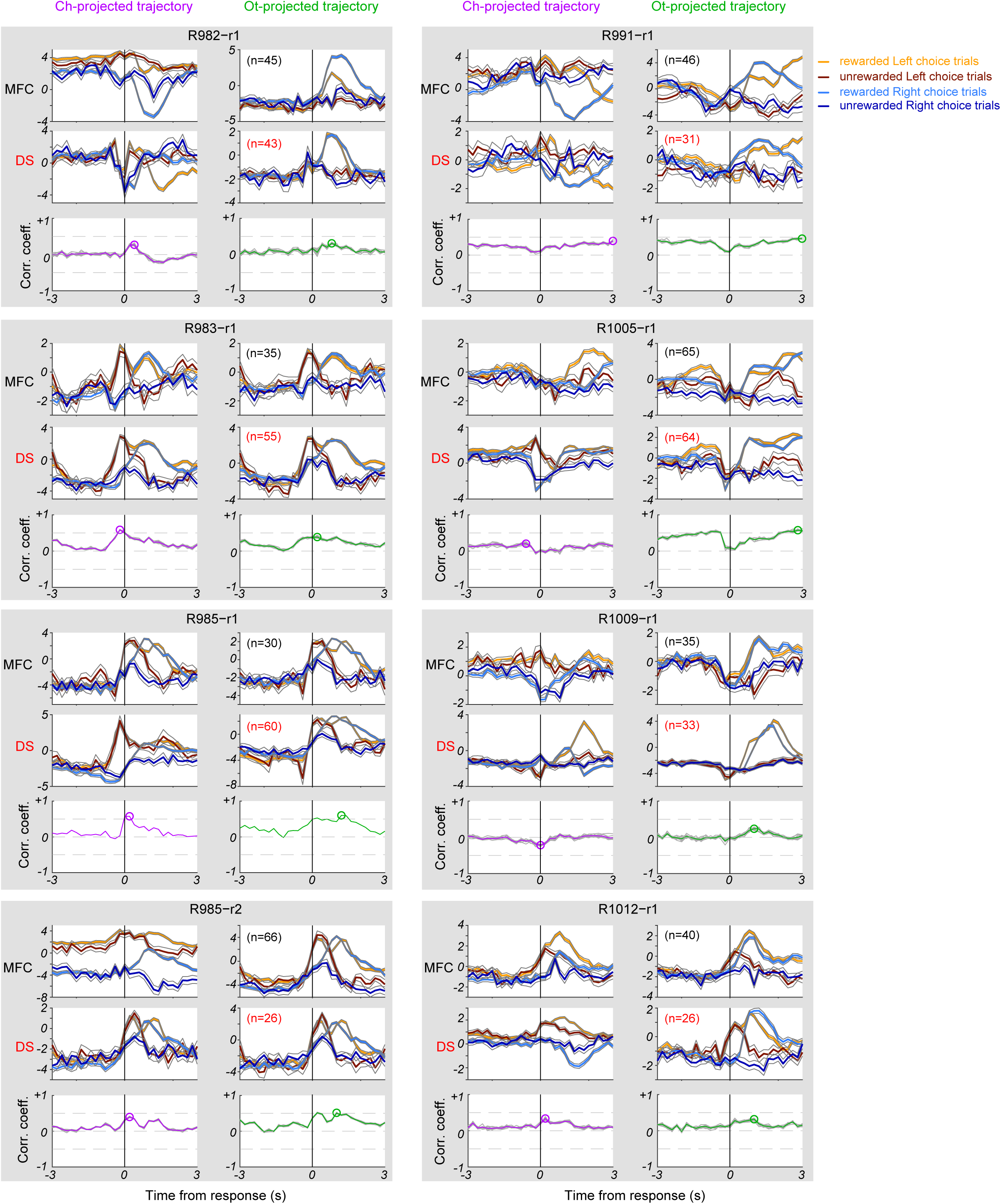
MFC and DS trajectories and their session-by-session correlations. Each gray box shows results from one session. Within a box, neural trajectory in MFC (*top*) and DS (*middle*) on choice (*left*) and outcome (*right*) axes are shown. Color codes indicate Left rewarded trials (orange), Left unrewarded trials (brown), Right rewarded trials (light blue) and Right unrewarded trials (dark blue). Thin lines represent S.E.M. (*Bottom*) Time evolution of correlation coefficient between MFC and DS trajectories. Gray lines show the correlation coefficients calculated with randomly sampled 431 trials in session R985-r1, which contains the minimum trial numbers among the 9 sessions. The averaged correlation coefficients between MFC and DS were displayed for Ch-projected (purple) or Ot-projected (green) trajectories. Circles indicate the times of maximum correlation.

For choice-selective or outcome-selective trajectories, the index *d’* peaked around the time of choice response or outcome, respectively (Figs. 4a and S3c). However, the peak times were not significantly different between MFC and DS (Fig. 4a). Then, we studied how neural trajectories evolved in each session by projecting the instantaneous rate vectors onto the axes orthogonal to choice (Ch) and outcome (Ot) hyperplanes obtained by the linear discriminant analysis (Fig. 4b). Hereafter, these trajectories are termed “Ch-projected” and “Ot-projected” trajectories. The temporal patterns of these trajectories varied from session to session in both MFC and DS (Fig. S4). However, their trial-averages exhibited transiently or gradually correlated increases between MFC and DS around choice (r = 0.62) and outcome (r = 0.50) events, respectively (Fig. 4b and 4e). The inter-trajectory correlations were particularly strong (Pearson correlation coefficient was about 0.5 or greater) in 4 sessions (R983-r2 in Fig. 4, R983-r1, R985-r1, and R985-r2 in Fig. S4). The neural trajectories diverged along with choice responses (Left and Right) around the response time (t1: 0.2 s before choice response), and afterwards diverged along with outcomes (reward and no-reward) during outcome period (t2: 1 s after choice response), as exemplified in Fig. 4c. Figure 4d shows the trial-by-trial values of Ch-projected and Ot-projected trajectories in MFC and DS at times t1 and t2, respectively. These trajectories were highly correlated between the two areas around the corresponding time points (Fig. 4e). In the other 5 sessions, while the Ot-projected trajectories were similar, the Ch-projected ones behaved differently in MFC and DS and the inter-trajectory correlations were also weaker between the two areas (Fig. S4).

**Figure 4.**
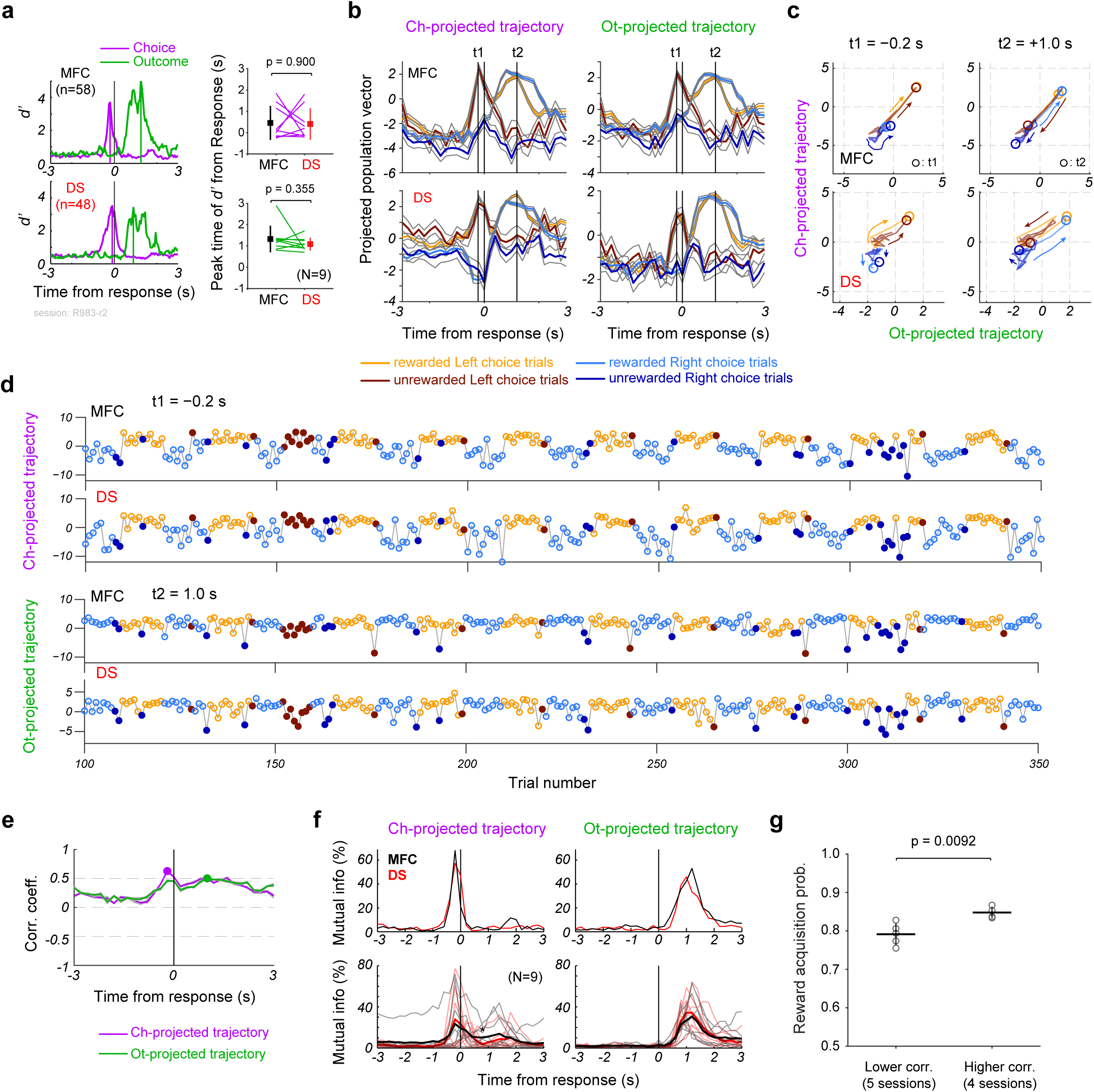
Temporal correlations of neural trajectories between MFC and DS. Unless otherwise stated, all results are shown for a typical session (R983-r2). (a) *Left*: Time series of discrimination degree (*d’*) for neural populations simultaneously recorded in MFC (n = 58, *Top*) and DS (n = 48, *Bottom*). Vertical lines indicate the peak times of *d’* measured between different choices (purple) and outcomes (green). *Right*: Comparison of the peak times of d’ between MFC and DS (N = 9 sessions) at Ch- and Ot-hyperplanes. Squares and error bars denote mean and SD. P-value was assessed by paired t-test. (b) Trial-averaged (*left*) choice-projected and (*right*) outcome-projected trajectories of population vectors in MFC (*top*) and DS (*bottom*). Gray lines show standard errors. Vertical lines indicate times t1 (−0.2 s) and t2 (+1.0 s) at which the trajectories were near maximally separated on choice and outcome axes, respectively. (c) Evolution of trial-averaged population vectors from -3 s to t1 (*left*) or t2 (*right*) in the space spanned by choice and outcome axes. Circles indicate population activities at t1 (*left*) and t2 (*right*). (d) The distances of choice- and outcome-projected trajectories from the separating hyperplanes at t1 (*top*) and t2 (*bottom*) are shown in MFC (*1st row*) and DS (*2nd* row). (e) Time evolution of Pearson’s correlation coefficients was calculated between MFC and DS projected trajectories. Purple and green lines represent the mean correlation coefficients, and gray lines show individual correlation coefficient calculated from randomly sampled 431 trials. Circles indicate the times of the maximum correlation. (f) *Top*: Mutual information about choice (*left*) and outcome (*right*) obtained from the trajectories in MFC (black) and DS (red) shown in b. *Bottom*: Mutual information from 9 sessions (thin) and the average (thick). Asterisk indicates statistical significance (Mann-Whitney U test, p <0.05). (g) Reward acquisition probability was significantly different between sessions with lower (< 0.5) and higher (> 0.5) trajectory correlations (t test, P <0.0092).

The enhanced correlations of Ch-projected and Ot-projected trajectories likely reflect choice responses and the monitoring of reward, respectively. As the degree of correlations varied during the task in all sessions, we addressed the behavioral variables correlated to the high MFC-DS correlation of neural trajectories. We investigated the temporal relationships among licking behavior, choice responses, reward delivery and the times of peak inter-trajectory correlations. As shown in Fig. 1e, the licking rate quickly reached a peak when the choice response was made and then decayed slowly during the outcome period (‘reward licking period’) in rewarded trials. In unrewarded trials, the decay was much faster. The peak correlation times of Ch-projected trajectories tended to cluster around choice response (t-test, p = 0.366) and reward delivery (t-test, p = 0.620), while those of Ot-projected trajectories were significantly later than choice response (t-test, p = 2.65×10^−3^) and reward delivery (t-test, p = 0.0297) (Fig. S5a), and were retained high during the following licking behavior (Fig. S5b). These results suggest that the epochs of highly correlated Ch- and Ot-projected trajectories are related to licking behavior or the intake of reward, respectively. The trajectory correlations exhibited different temporal profiles in rewarded and unrewarded trials (Fig. S5b), but the peak correlations of Ch- (Wilcoxon signed-rank test, p = 0.218) and Ot-projected (p = 0.546) trajectories were not significantly different between the two trial types, suggesting that the enhanced correlations reflect the monitoring of reward rather than actual reward delivery.

**Figure S5.**
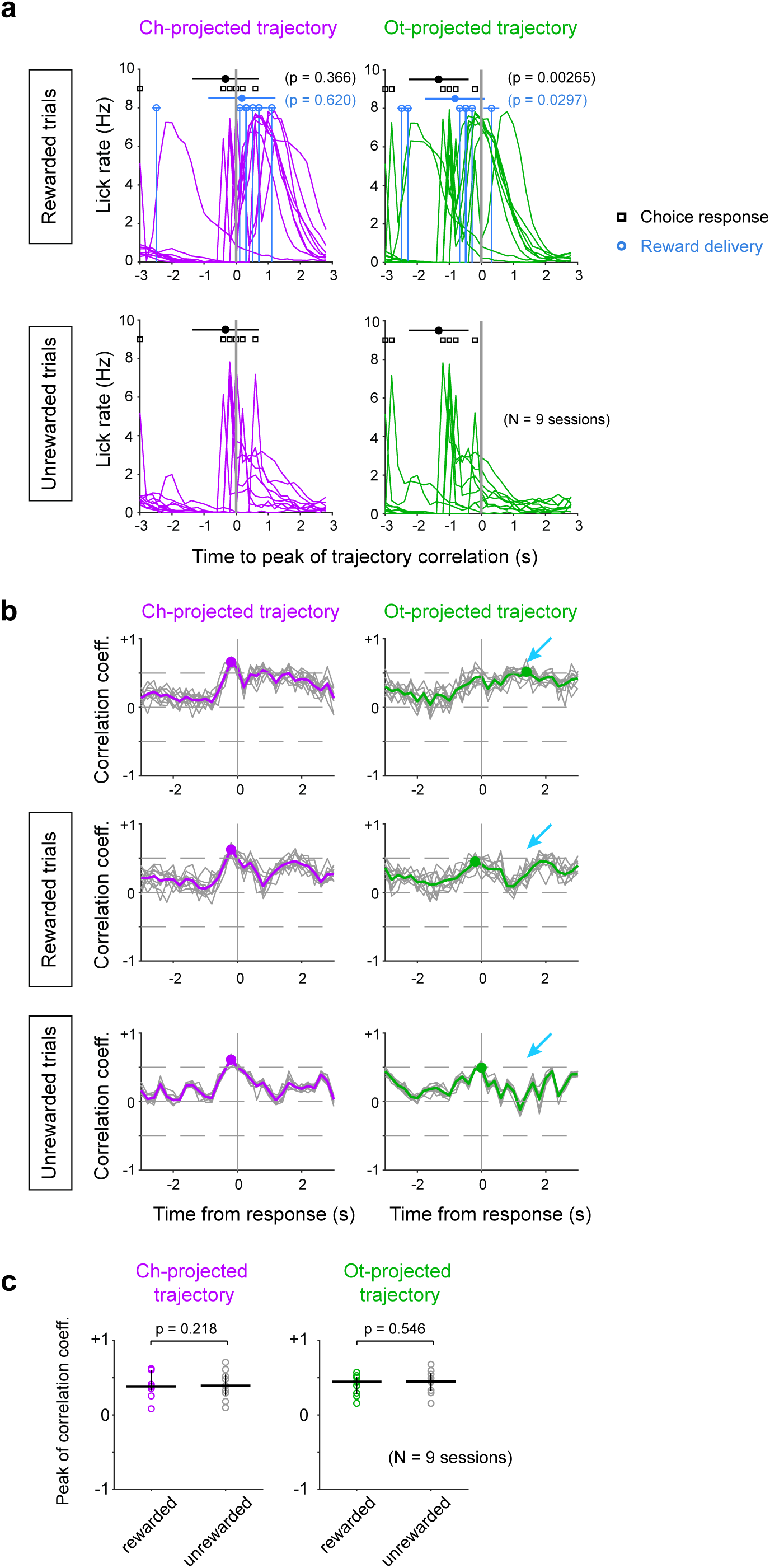
Behavioral variable when the correlation of trajectories reached peak. (a) Session-by-session licking rate, choice response (black) and reward delivery (light blue) are aligned at the time when Ch-projected (purple) and Ot-projected (green) trajectories showed peak correlations between MFC and DS. The origin of time refers to the time of peak correlations. Means (filled symbols) and SDs (horizontal lines) of choice response and reward delivery time are shown (N = 9 sessions). (b) Time evolution of trajectory correlations are shown for a rat in rewarded (middle), unrewarded (bottom), and mixed (top) trials. Arrows indicate the times of peak correlation in the mixed trial condition. We randomly selected 63 trials in each condition and calculated correlation coefficient (gray line). We repeated this procedure by 10 times. Averaged correlation is represented by colored thick line. Circles indicate the peak values and times. (c) Comparison of peak correlation coefficients between rewarded and unrewarded trials.

It is intriguing to examine whether neural trajectory switched in either or both of MFC and DS when the rats switched their choices after unrewarded trials. In the present task, rats switched their choices after a reversal of reward block within a couple of unrewarded trials (Fig. 1b and c). We hypothesized that during the unrewarded trials neural trajectory should cross the hyperplane (Unless otherwise stated, hyperplane refers to Ch-hyperplane) from one side to the opposite side (i.e. ‘Right to Left’ or ‘Left to Right’) before a choice response in the next trial. Out of 9 sessions, in 3 sessions we observed a gradual crossing of MFC trajectory during ITIs, while a similar crossing of DS trajectory was observed only in one session (Fig. S6). Although the crossing was somewhat more prominent in MFC than in DS, it was difficult to determine which region dominated switching choice responses.

We further verified the analogous encoding of choice and outcome information in MFC and DS by calculating the time evolution of the information carried by the neural trajectories (Fig. 4f). The amount of information varied from session to session and was slightly greater in MFC than in DS. However, the mutual information averaged over nine sessions was not statistically different between MFC and DS at most of the times around choice response (Mann-Whitney U test, p >0.05). These results indicate that neural trajectories evolve in parallel in MFC and DS and transiently increase the mutual correlation to transfer choice and outcome information without a large loss. Reward acquisition probability was significantly larger (p = 0.0092, two-sample t test) in sessions with high inter-trajectory correlations (> 0.5) during choice and outcome periods than in sessions with low correlations (Fig. 4g). In this analysis, the session R1005-r1 was included in the low-correlation group because the correlation was low during choice and outcome periods. Omission of this session did not change the essential result of comparison (p = 0.0193, two-sample t test).

**Figure S6.**
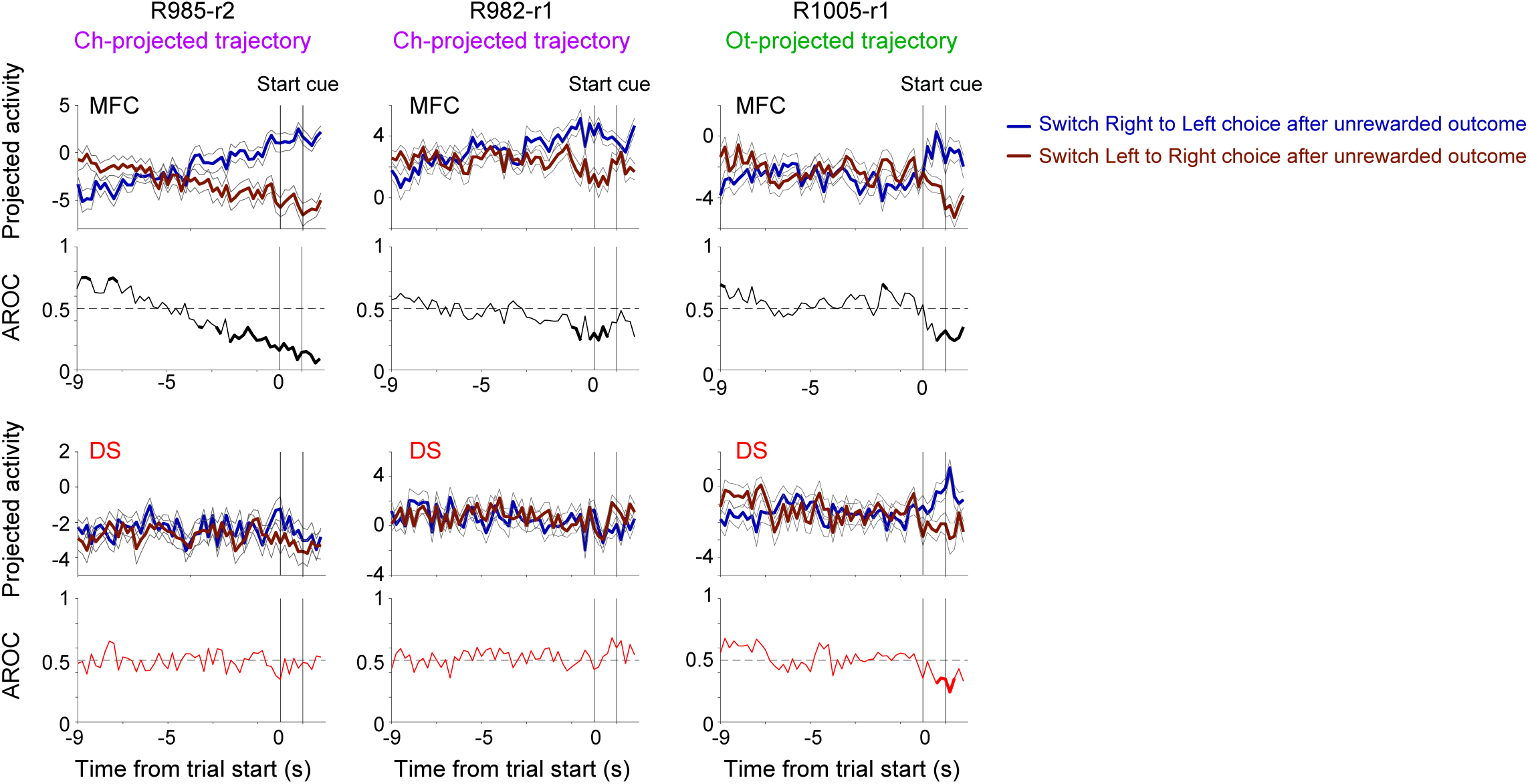
Crossing trajectories between two switching choice trials during inter-trial interval (ITI) period. Neural trajectory during ITI period post unrewarded episode in 3 sessions (left: R985-r2, Ch-projected trajectory; middle: R982-r1, Ch-projected trajectory; right: R1005-r1, Ot-projected trajectory) in MFC (1st row) and DS (3rd row). Dark blue and brown denote the neural trajectories when rats switched choices from Right to Left and from Left to Right, respectively. Black thin lines denote standard errors. Differences in the two trajectories were quantified by AROC values in MFC (2nd row, black) and DS (4th row, red). A gradual shift from high values (> 0.5) to low values (< 0.5), suggests that the trajectories evolved to the opposite side of the corresponding hyperplane. Statistical significance of AROC values is estimated by Mann-Whitney U test (p <0.05) and represented by thick lines. Two vertical lines show the duration of start cue presentation.

### Synchronous spiking when neural trajectories were highly correlated between MFC and DS

As shown above, neural trajectories in MFC and DS exhibit task event-dependent modulations of coherence. This result raises questions about whether neuronal firing is also temporally correlated between these regions and whether correlated spikes, if any, are related to neural trajectories. To study these questions, we calculated cross-correlograms (CCGs) of MFC-DS cell pairs in the time window in which MFC-DS correlations of trajectories were maximized (see Fig.4e and Fig. S4) (Methods). Then, we compared task-event relevant and irrelevant CCGs in three trial categories (i.e., Left-choice, Right-choice and rewarded trials).

Our analysis revealed MFC-DS cell pairs displaying statistically significant spike synchrony in a task-event-dependent manner (Fig. 5a). Shuffling spike trains across trials within the individual pairs eliminated the peaks in CCGs (Fig. S7), implying that these cell pairs fired with a close temporal relationship in each trial. The peak values of CCGs showed significant correlations between MFC and DS (> 4 SD of the base line) in task-event related windows, but these values were greatly decreased if CCGs were calculated in task-event-irrelevant windows (i.e., at random time points) (Fig. 5a and b). These results suggest that the emergence of spike synchrony between MFC and DS cells is correlated with task events.

**Figure 5.**
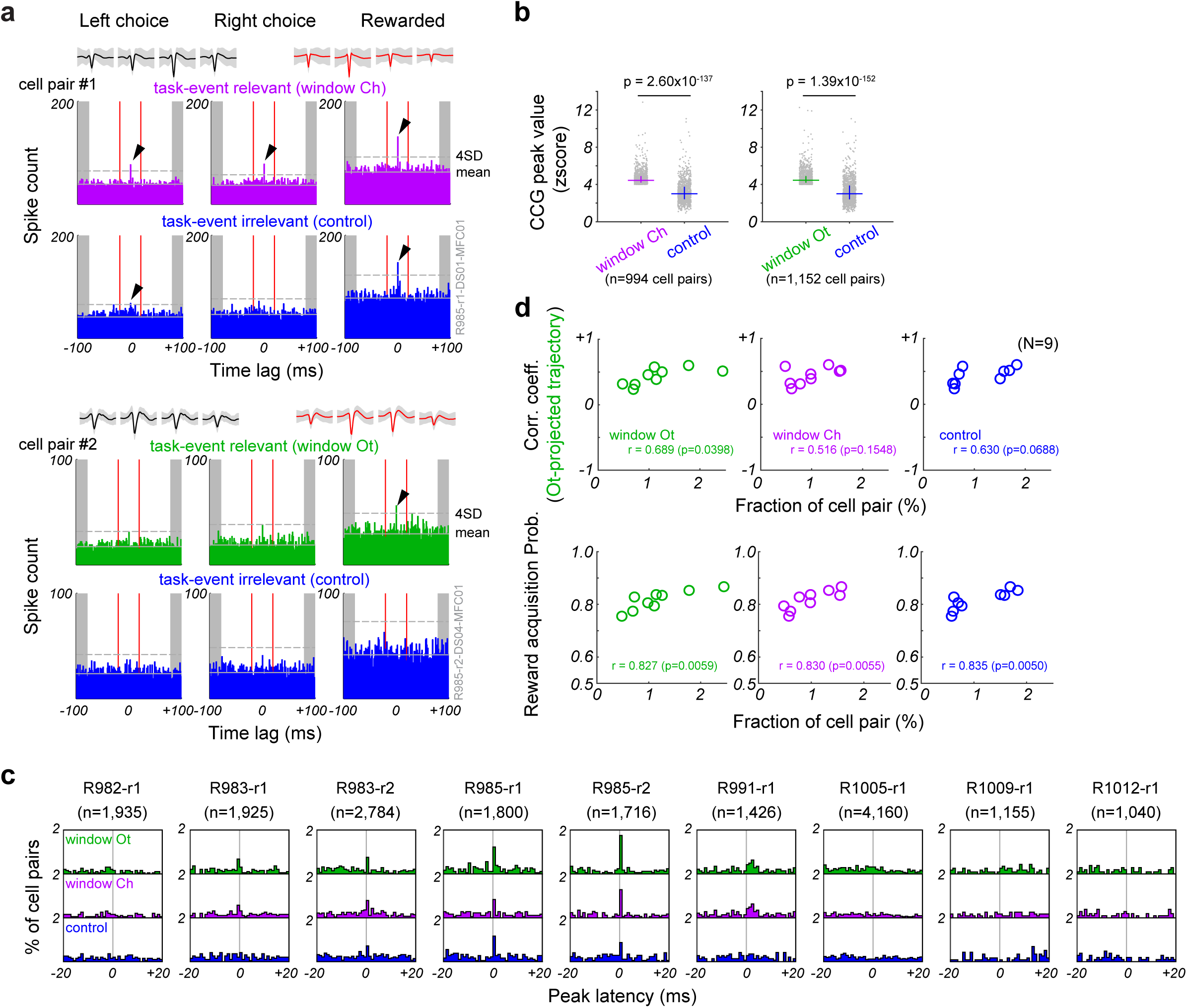
Relationships among MFC-DS spike synchrony, trajectory correlation and task performance. (a) Spike CCGs of two MFC-DS cell pairs. For each cell pair, time lags of MFC cell spike firing relative to DS cell spike firing were calculated in task-event relevant (*middle row*) and irrelevant (*bottom row*) time windows. Average and SD (gray shade) of spike waveforms recorded with tetrodes in MFC (black) and DS (red) are also shown (*top row*). *Left and middle columns*: Left or Right choice trials with arbitrary outcomes, respectively. *Right column*: rewarded trials with arbitrary choice positions. Windows Ch (purple) and Ot (green) refer to the analyzed time windows centered around the peak MFC-DS correlations of Ch-projected and Ot-projected trajectories, respectively. Task-event irrelevant CCG gives a control and was calculated at a randomly selected time point. Arrowhead indicates the peak of CCGs above the mean + 4SD of the baseline defined by spike counts from -100 to –80 ms and from +80 to +100 ms (gray shaded ranges). (b) CCG peak values of cell pairs in windows Ch (purple) and Ot (green) are compared with those in task-event irrelevant windows (black). Each dot represents a cell pair with a significant CCG peak (peak > 4SD). Horizontal lines and error bars show the median and 1^st^/3^rd^ quartiles, respectively. P-values were calculated by Wilcoxon signed-rank test. (c) Proportion of MFC-DS cell pairs that displayed an excess amount of spike coincidences in rewarded trials was plotted for the 9 sessions against CCG peak time lags. The numbers of such cell pairs are given in parentheses. (d) (*Top*) Correlation coefficients between MFC and DS Ot-projected trajectories are plotted against the fraction of MFC-DS cell pairs showing a CCG peak within ±2 ms for different time windows. (*Bottom*) Relationship between such MFC-DS cell pairs and animal’s reward acquisition probability. Symbols represent individual sessions, and Pearson correlation coefficient and P-value are denoted for each diagram.

Spike synchrony tended to occur when neural trajectories were highly correlated between MFC and DS, which further supports the role of spike synchrony in coordinating neural population activities in the two regions. The spike count of MFC cells exhibited statistically significant peaks at near zero time lags from spikes of DS cells in 4 sessions (R983-r1, R983-r2, R985-r1 and R985-r2 in Fig. 5c). At this epoch, the maximum correlation coefficients between MFC and DS trajectories were near or over 0.5 (Fig. 4e and Fig. S4). By contrast, the CCG distributions showed no significant peaks in the other 5 sessions (R982-r1, R991-r1, R1005-r1, R1009-r1, and R1012-r1). However, in these 5 sessions the correlations of neural trajectories were also weaker compared to the former 4 sessions (Fig. S4). Interestingly, in rewarded trials the correlation coefficient between Ot-projected trajectories was correlated with the fraction of cell pairs showing CCG peaks within ±2 ms (Pearson correlation test, window Ot, r = 0.689, p = 0.0398) (Fig. 5d, top). Further, the fraction of such cell pairs was correlated with the probability of animal’s reward acquisition (Fig. 5d, bottom). These results suggest that precisely correlated firing of MFC and DS cells underlies the coherent evolution of neural trajectories between these areas and boost animal’s task performance.

**Figure S7.**
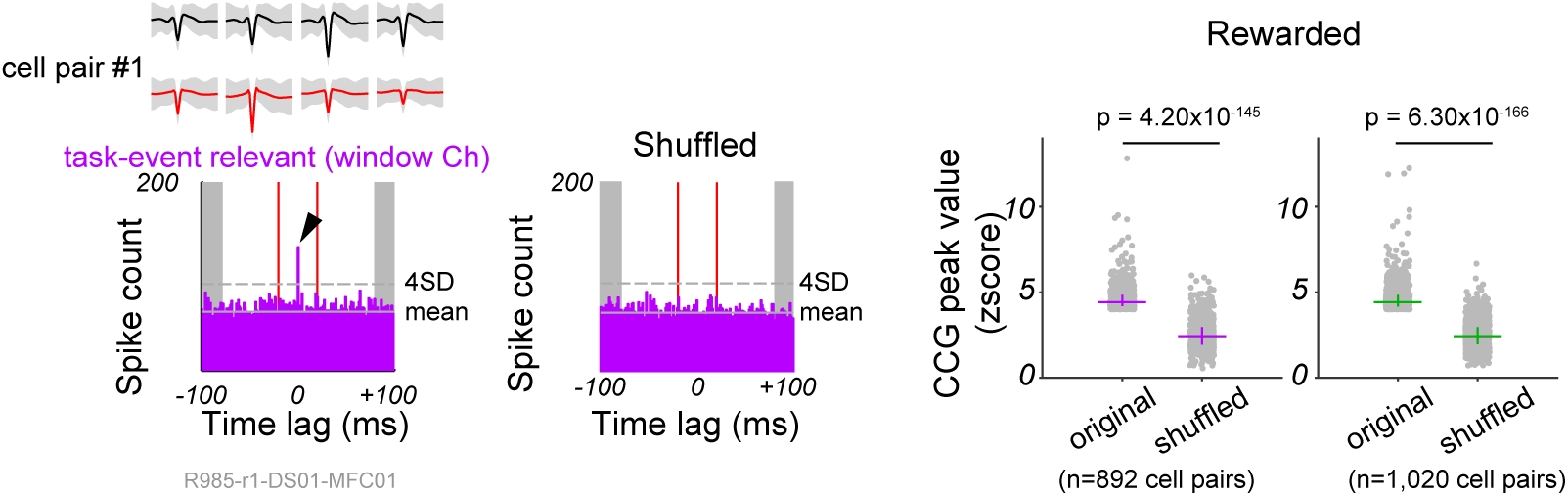
Significant cross correlations between MFC and DS cell were dependent on the spike timing in the ongoing trial but not on the number of spikes. Left: CCGs of the cell pair #1 in Fig. 5a were calculated in the original (left) and shuffled (right) trial orders in rewarded trials. Right: Normalized CCG peak amplitudes were calculated for all cell pairs showing significant CCG peaks in windows Ch (purple) and Ot (green). The peak amplitude are compared with those obtained in shuffled trial orders. Horizontal lines and error bars show the median and 1st/3rd quartiles, respectively. P-values were calculated by Wilcoxon signed-rank test.

### Inactivation of MFC attenuated neural representation of choice and outcome in DS cells

To confirm the contributions of inter-trajectory correlations between MFC and DS to adaptive control, we examined the effect of bilateral MFC inhibition on the choice behavior and DS-cell activity in 5 rats (muscimol group) (Fig. 6a, Methods). It is known that MFC is one of the main regions projecting excitatory inputs to DS [6-8]. Another 10 rats were examined the choice behavior without bilateral MFC inhibition as a control group. The muscimol injection induced variable choice patterns in five muscimol-treated rats (F test, F = 16.95, p = 0.0179). After the injection (Day 2), the reward acquisition probabilities of four rats were significantly different from those prior to the injection (Day 1) (χ2 test, p<0.05). Among these four, the reward acquisition behavior was impaired in three while another rat rather improved the behavioral performance (Fig. 6b). Figure 6c shows an example of impaired task performance. This rat reduced “stay” behavior after preceding rewarded outcomes but increased such behavior after preceding unrewarded outcomes (Fig. 6c-d). Indeed, the fractions of “stay” and “switch” choice patterns were significantly different between Day1 and Day2 in the muscimol group (two-way ANOVA, F_1,4_ = 5.774, p = 0.021) but not in the control group (F_1,9_ = 0.248, p = 0.620). Accordingly, the control and muscimol groups showed significantly different reward acquisition probabilities between Day 1 and Day2 (two-sample t-test, p = 0.0472). The behavioral variability between the two groups was unlikely attributed to the attenuation of their motivations for the task because the number of trials was not significantly different between Day1 (mean ±SD = 731 ±86 trials) and Day2 (740 ±96 trials) in the muscimol group (unequal variance t-test, p = 0.877) and these numbers were not significantly different from those of the control group. Those results suggest that MFC is not a sole contributor to the present adaptive choice behavior.

**Figure 6.**
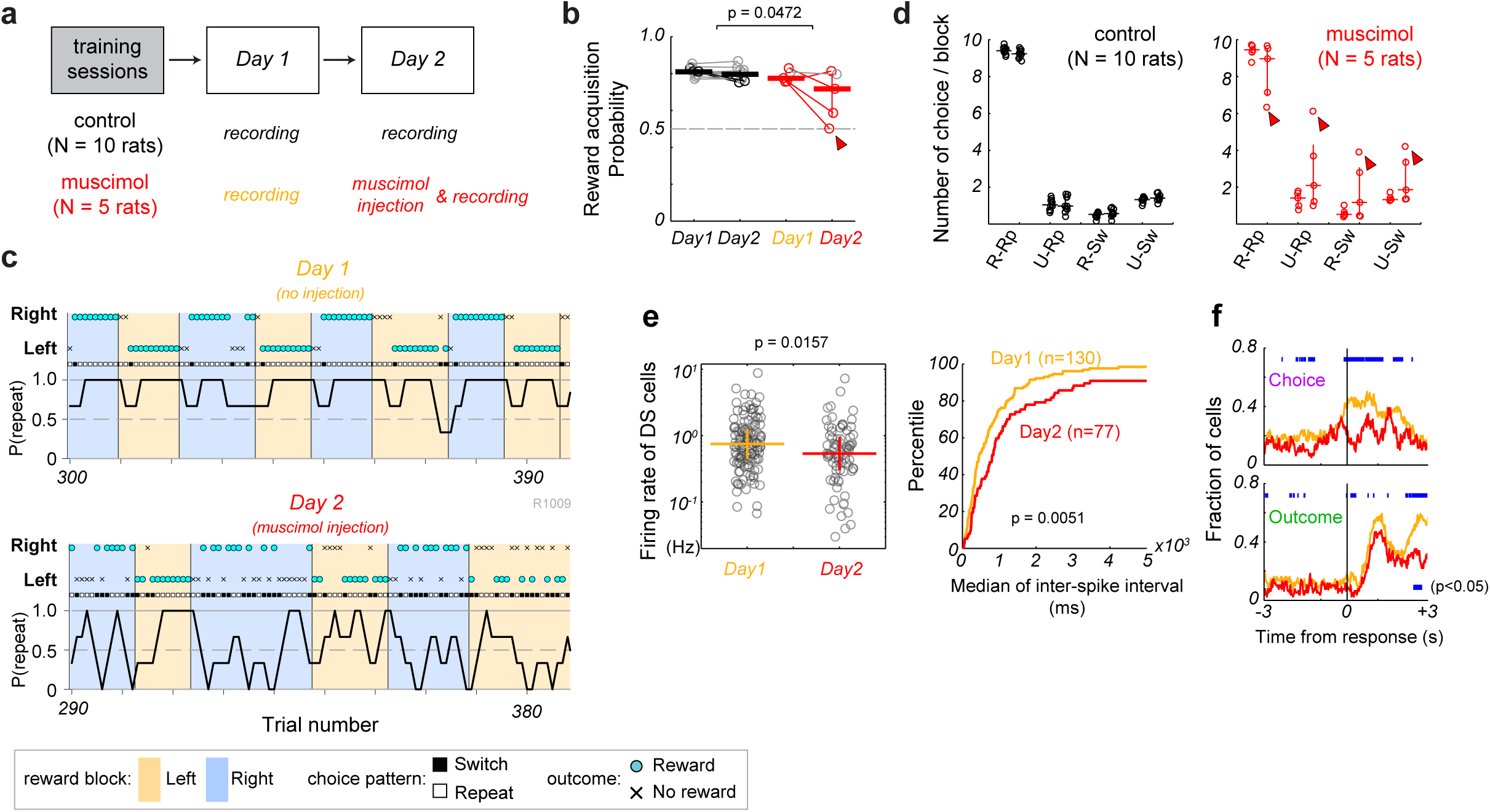
Inactivation of MFC attenuated neuronal representation of choice and outcome in DS. In statistical comparisons, horizontal and vertical lines indicate the median and 1st/3rd quartiles, respectively. (a) Experimental schedules for control (N = 10 rats) and muscimol (N = 5 rats) groups. At the first session (Day1), all 15 rats underwent the recordings of multi-neuron activity from MFC and DS without any pharmacological treatments. At the second session (Day2), we injected muscimol into the MFC of five rats (muscimol group) and recorded multi-neuron activity from DS. For the remaining 10 rats (control group), neural activities were recorded from MFC and DS without any injection. (b) Reward acquisition probabilities in Day1 and Day2 are shown in control (black) and muscimol (red) groups. Each symbol shows data from a rat. Colored (black or red) symbols indicate the individual rats that showed significantly different probabilities (χ2 test, p <0.05) between Day1 and Day2. Arrowhead indicates the rat shown in c. The differences in reward acquisition probabilities between Day1 and Day2 were compared between groups by two-sample t-test. (c) An example of impaired adaptive choice behavior of a muscimol-injected rat is shown on Day1 (top, no injection) and Day2 (bottom, muscimol injection). (d) Mean number of choice patterns per block on Day1 (left) and Day2 (right). Left: control group, Right: muscimol group. R: rewarded in last trial, U: unrewarded in last trial, Rp: repeat in current trial, Sw: switch in current trial. (e) Firing patterns of DS cells were compared (Mann-Whitney U test) in the muscimol group in terms of firing rate (Left) and inter-spike interval (Right) between Day1 (n = 130, orange) and Day2 (n = 77, red). Circles indicate the firing rates of individual DS neurons. (f) Time evolution of the fraction of (top) choice-modulated and (bottom) outcome-modulated DS cells (two-way ANOVA, 200-ms-long sliding-by-20-ms bins, p <0.05) on Day1 (orange) and Day2 (red). Blue bars show the epochs of significantly different fractions between Day1 and Day2 (χ2 test, p <0.05).

The inhibition of MFC diminished neuronal activities and their selective modulations in DS. The firing rates of DS cells (n = 77) were significantly lower (Mann-Whitney U-test, p = 0.0157) and the median of inter-spike intervals was significantly longer (Mann-Whitney U-test, p = 0.0051) on Day2 than on Day1 (n = 130) (Fig. 6e), suggesting an attenuated excitatory drive or an enhanced inhibition on DS cells. The MFC inhibition also altered the task-related activities of DS cells, especially those encoding choices and outcomes (two-way ANOVA, p <0.05). The proportion of choice-modulated cells around and after choice responses was significantly smaller on Day2 than on Day1 (χ2 test, p <0.05) (Fig. 6f, top), and so was the proportion of outcome-modulated cells during the latter part of outcome period (more than 2 s after a choice response). However, this proportion was not significantly different during the former part of outcome period (Fig. 6f, bottom). Taken together, our results suggest that neuronal activation in MFC is engaged in DS neural coding both choice-related information around response time and outcome-related information during reward licking behavior.

## Discussion

This study revealed a highly parallel evolution of MFC and DS neural trajectories regulating outcome-based choice behavior. These trajectories initially encoded a current choice and later reflected the outcome of the choice response. Despite variation in temporal patterns of the trajectories across sessions and individuals, the amount of information carried by the trajectories was not significantly different between MFC and DS. Furthermore, cross-area spike coincidences in the millisecond range were enhanced in rewarded trials when the trajectories were highly correlated between the two regions, suggesting that the parallel trajectory evolution in MFC and DS actively participates in coordinating the behavioral task.

Along the frontal cortex-basal ganglia axis, the MFC-DS channel is thought to be the stage to process action selection based on past outcomes [1,3]. A widely hypothesized mechanism for this process is that neural ensemble in DS selectively gates the necessary information (or suppresses the unnecessary information) received from MFC. However, our results provide no support for this hypothesis because neural populations in MFC and DS preserve similar amounts of information about choices as well as outcomes and their trajectories evolve quite similarly in the two regions (Fig. 4).

Our data suggest in contrast to the gating hypothesis that the neural trajectory of DS ensembles reflects that of MFC ensembles, which is consistent with previous work demonstrating a simultaneous and correlated activation of time-dependent ramping activity in MFC and DS during a temporal judgment task [36]. An alternative interpretation, given that MFC innervates DS unidirectionally, is that MFC cells exhibit a choice-related signal prior to DS cells and then convey the signal to DS in a sequence, as previously suggested [14,37]. However, our data does not support this interpretation because choice-related signals concomitantly emerge in MFC and DS. This discrepancy may be attributed to several differences in experimental conditions. For instance, unlike other studies, we recorded neuronal activities in a head-restrained condition, did not change the probability of reward delivery during the task, and directly compared simultaneous neuronal activations between MFC and DS.

A possible role of the corticostriatal circuit suggested by our findings is that construction of temporal correlation of neural activation between MFC and DS could lead to consolidating efficient transmission of information from frontal cortex to downstream of basal-ganglia or to Inhibiting inappropriate action. A previous study demonstrated the synchronization of oscillatory local field potentials at 5-13 Hz across motor cortex, DS and substantia nigra pars reticulata, which is the output terminal of the basal ganglia, and the propagation of spindle-like spike-and-wave oscillations along this cortico-basal ganglia pathway in freely moving rats. Spikes of individual neurons were phase-locked to the oscillatory local field potential and synchronized within the cortical-basal ganglia network [38]. Since synchronized neuronal discharges may enable a reliable information transmission [29,30], the temporal correlation between neural ensembles in the frontal cortex-basal ganglia network could be a consequence of learning the decision behavior. In accordance with this view, during skill learning, neural ensembles in both motor cortex and DS are known to develop spike correlations [39,40] and precise temporal spiking patterns [31,41]. In the present study, synchronous spiking was coordinated between MFC and DS cells when neural trajectories in these areas were strongly correlated in the well learned rats, supporting the hypothesis (Fig. 5).

These results raise the question about how such a precise spike synchrony, which is indicated by the short latency of CCG peak between MFC and DS cells (Fig. 5), was generated between the cortical and subcortical regions. There are at least two possible explanations. One possibility is that corticostriatal neurons in MFC make monosynaptic connections to the postsynaptic DS cells. In this case, presynaptic MFC cells should discharge prior to the firing of postsynaptic DS cells, but this was not the case at all in the CCGs between these cells (Fig. 5). Another possibility is that the firing of MFC and DS cells was driven by a common input from other corticostriatal neurons in the MFC [42,43] or neurons in other cortical areas projecting to both MFC and DS. Medium spiny DS cells receive excitatory inputs from ipsilateral and contralateral MFC [16], while MFC cells’ output projects to both ipsilateral and contralateral cortical areas and DS through corpus callosum, the so-called crossed-corticostriatal (or intratelencephalic) cells [6,44]. Since excitatory connections are abundant between MFC corticostriatal pyramidal cells [45], synchronous discharges of these neurons generate large EPSPs in their postsynaptic target cells in MFC and DS, which increases the probability of synchronous firing between MFC and DS cells. Alternatively, both MFC [46] and DS [7,8] receive afferent connections from higher-order cortical areas such as the ventrolateral OFC and PPC. Since the OFC and PPC are implicated in goal-directed decision-making [47,48], excitatory inputs from a common subpopulation of OFC and/or PPC neurons may evoke synchronous spiking of MFC and DS cells during decision-making. In the present study, bilateral MFC inactivation attenuated, but did not completely eliminate, choice and outcome representations in DS cells (Fig. 6), suggesting that brain regions other than MFC also drive DS cells in outcome-based action selection. For instance, optogenetic inhibition of PPC corticostriatal projection neurons altered the history dependent choice bias during decision making [49]. To distinguish between the different causes of spike synchrony between MFC and DS cells further recordings from multiple regions involving MFC, OFC, PPC, and DS are required.

It was recently shown in the three-layered cortex of the reptile that just a few spikes of single pyramidal neurons can trigger a cascade of firing sequences of neuron ensembles [50], as was suggested by theoretical studies [29,51]. Our results indicate the possibility that a similar cascade of firing sequences is triggered in the cortico-basal ganglia pathway during the parallel evolution of frontal cortical and striatal neural trajectories. Such spike sequences may not only enhance reliable information transmission (i.e., spikes) from the cortex to the basal ganglia, but also may contribute to regulating the gain of synaptic plasticity together with modulations by dopaminergic afferents [52,53]. Further experiments are required to clarify these points.

In sum, we have shown that neural trajectories evolve in DS during goal-directed decision making in parallel with a similar trajectory evolution in MFC. When the parallel trajectory evolution in DS and MFC exhibits a strong coherence, spike synchrony is also enhanced between the two regions and behavioral performance tends to be improved. Our results suggest that the parallel evolution of cortical and striatal neural trajectories is a hallmark of successful decision making.

## Acknowledgments

We thank T. Sharp and T. Takekawa for their advice and supports for initial analysis on our data. We are grateful to M. Gilson, P. Goncalves, J. Igarashi, Y. Isomura, T. Kurikawa, and M. Tatsuno for their valuable comments on our results. We thank J. Wickens for his critical reading of our manuscript and Y. Goda for her advices. This work was partly supported by the Grants-in-Aid for Scientific Research (KAKENHI) from MEXT (no. 24700345 to TH, nos. 18H05213 and 19H04994 to TF).

## Author contributions

TH and TF designed the project. TH and RH conducted experiments. TH analyzed data. TH and TF wrote the manuscript.

## Methods

### Animal preparation

All experiments were approved and carried out in accordance with the Animal Experiment Plan by the Animal Experiment Committee of RIKEN. Male Long-Evans rats (N = 34, 6 weeks, 200-220 g, Japan SLC, Inc.) were used. Home cages were set in a temperature and humidity controlled environment with lights maintained on a 12-h light/dark cycle. Prior to a primary surgery, rats were handled briefly and habituated to a stainless-steel cylinder in the cages. All surgical procedures were operated under sterile circumstances. Animals underwent three surgical procedures. Rats were anaesthetized with 2% isoflurane. Their body temperature was monitored with a rectal probe and maintained at ∼37°C on a heating pad during the surgery.

At a primary surgery, a sliding head-attachment (Narishige) was implanted on the skull with dual-curing resin cement (Panavia, Kuraray Noritake Dental) and dental resin (Unifast II, GC) as done before in our laboratory [12,13,23,54]. Reference and grounding electrodes (teflon coated silver wire) were put on dura mater above the cerebellum. After recovery from the surgery, the rats were deprived water intake in the home cages in order to utilize water as a reward for the execution of a task, although food was available *ad libitum*. Water (10 ml) was supplemented at every weekend. At a second surgery 3 days prior to a first recording session, we injected a retrograde tracer Fluoro-Gold (FG) into the DS in order to confirm if our recording sites corresponded to the MFC sending corticostriatal projection to the DS. A glass micropipette filled with 2% FG (Fluorochrome) dissolved in 0.1 M cacodylic acid was installed on a micromanipulator angled medially by 27 deg. The pipette was inserted through a tiny burr hole drilled in the skull over left hemisphere (+1.5 mm to Bregma, 1.0 mm lateral to midline, 4.3 mm traveling distance) so that the pipette tip reached the dorsocentral part of striatum (+1.5 mm anterior to Bregma, approximately 3.0 mm lateral to midline, approximately 3.8 mm ventral to pia mater), which rostral agranular medial cortex (AGm) pyramidal neurons directly innervate (Figs. 2 and S1). FG was iontophoretically loaded through 7 s pulse of +5.0 μA by 7 s interval for 30 or 60 minutes with an iontophoresis pump (BAB-501, Kation Scientific). Then, the tiny hole was covered with sterilized spongel and dental silicone sealants (Matsukaze). At a third surgery after the training sessions were completed, two cranial windows (1.2 mm diameter) were made above the DS and MFC of left hemisphere (+1.0 and +3.0 mm to Bregma, 3.0 and 1.0 mm to midline for DS and rostral MFC, respectively) and then its dura maters were removed. For bilateral muscimol injection session, another tiny burr hole was additionally drilled in the skull above MFC of right hemisphere (+3.0 mm to Bregma, 1.0 mm to midline). The cranial windows were covered with the silicone sealants.

### Behavioral training apparatus

After recovery from the first surgery, the rats were trained to perform a behavioral task controlled under a customized multiple-rats training system (O’hara), which enabled training several rats to learn a task paradigm in parallel [23,54]. The behavioral task was controlled by a custom-written software with LabVIEW (National Instruments). In an isolated training chamber, each rat was placed at a body-supporting cylinder, and its head was fixed rigidly and painlessly by screwing a sliding head-holder on a stereotaxic frame. Auditory stimuli were presented at 60-dB SPL via a speaker placed in front of the rat. The timing and direction of licking movement were detected when its tongue interrupted an infrared beam placed below the left and right spouts. The detection sensitivity was much enough to detect tongue movements during licking spout, but not sensitive enough to detect other potential movements such as whisking and mastication. White plastic plates were located at both besides of the face in order to prohibit rats from approaching the beam detectable spaces by forelimb reaching (Fig. 1a). Spouts were connected to a syringe set on a single-syringe pump (AL-1000, World Precision Instruments) via silicon tubing. Water delivery from each spout was regulated by an audible pinch valve triggered by TTL signal which also triggered the syringe pumping.

### Behavioral task

A trial began with a pure tone presentation (3 kHz, 1 s) (Start in Fig. 1a). Rats were required not to lick any spouts from 0.3 s after Start cue offset until an appearance of another auditory cue (10 kHz, 0.2 s) (Go). If the rats licked any spouts during the delay period, the trial was immediately aborted. This pseudorandom delay period ranged between 0.7-2.3 s. After the Go signal onset, the rats were allowed to lick one of either left or right spout within a response window (5 s). A first lick was judged as a choice response (Choice). When the chosen spout location corresponded to ongoing reward location, 0.1% saccharin water (15 μl) was delivered as a reward after a pseudorandom delay period ranging between 0.3 and 0.7 s (2nd Delay). After an outcome period (4 s, Outcome), a next trial began. On the other hand, when the rats chose no-reward spout, they did not receive any sensory feedback but had an additional time-out of 5 s after the outcome period. After the time-out, a next trial began.

At a first training session, the rats learned to associate licking a single spout located at left (or right) side with a reward delivery. Once the rats voluntarily licked the spout to acquire a reward, they learned to associate licking another spout set at the opposite side with a reward acquisition through another spout. At this stage, the second delay period after choice was fixed as zero (no delay). At a second training session, the reward-associated spout location was switched after a bunch of cumulative rewarded trials were observed at each side. As rats reliably showed repetitive rewarded choices rather than random choices, the number of reversal of reward-location increased. Eventually, once the cumulative rewarded trial number reached 10 within each block, the reward-associated spout position was reversed without any feedback like sensory or physical differences in the task. Therefore, rats could not know the block reversal *a priori* without experiencing forthcoming trials. From 13th and 17th sessions, the first and second delay periods were prolonged by extending to 0.7-2.3 s and to 0.3-0.7 s, respectively. After each rat was trained at a training chamber for 19 days, they were afterwards habituated to another task chamber, where electrophysiological recording experiments were conducted, through two more training sessions.

### Electrophysiological recordings

Two daily recording experiments were conducted. Multi-neuron activity was simultaneously recorded from MFC and DS of left hemisphere with two 32-channels silicon probes consisting of four shanks (0.4 mm shank separation), on which tetrode-like electrode sites were spaced vertically by 0.5 mm (A4×2-tet-7/5mm-500-400-312, NeuroNexus Technologies). Each probe was connected to a custom-made headstage installed on either one of two fine micromanipulators (1760-61; David Kopf Instruments) mounted on a stereotaxic frame (SR-8N, Narishige). A silicon probe was penetrated vertically (depth from pia mater: 1.2 mm) into MFC (+2.4-4.2 mm to Bregma, 1.0-1.4 mm to midline), and the shanks were aligned along midline. Another silicon probe angled posteriorly by 6 deg was inserted into DS through a cranial window (+0.6-1.0 mm to Bregma, 2.1-3.7 mm to midline, 4.0 mm traveling distance), and the shanks were aligned along coronal suture. Multiunit signals were amplified by the headstages before being fed into main amplifiers (Nihon Koden, 2000 gain) with a band-pass filter (0.5 Hz to 10 kHz). At the second recording session, juxtacellular activity was recorded from the MFC of left hemisphere. A glass microelectrode was prepared by a laser puller (P-2000; Sutter Instrument) and filled with 2-3% Neurobiotin (Vector Laboratories) dissolved in 0.5 M potassium chloride (9-19 MΩ). The electrode was inserted into MFC through the cranial hole for MFC silicon probe with a stereotaxic hydraulic micromanipulator (SM-25C, Narishige). Juxtacellular activity was amplified (final gain 1000) with two amplifiers (IR-283, Cygnus Technologies; EX4-400, Dagan) and filtered (0.3-10 kHz). All neural data was sampled at 20 kHz with two hard-disc recorders (LX-120, TEAC), together with time of task events and licking each spout (left and right). After juxtacellular recording, we tried to electroporate Neurobiotin into the recorded cell with positive current injection (2-14 nA, duration of 0.5s at 1 Hz interval, for 5-15 mins) in order to visualize the recorded cell *post hoc* and to verify the recording location.

### Inactivation of the MFC

Of 34 rats, 15 reached a performance criterion (>75%) at the first recording session. Of the 15 rats, 10 and 5 were tested without and with injection of GABAa agonist muscimol into bilateral MFC at the second recording session, respectively. For the muscimol group, the rats were placed in the body restrainer and their heads were fixed on the stereotaxic frame. A microinjection syringe (Hamilton) filled with 0.1% muscimol (muscimol hydrobromide, Sigma-Aldrich) dissolved in 0.1M phosphate-buffered saline [55] was set on a microsyringe pump (Legato 130, KD Scientific) which was installed on the micromanipulator (1760-61; David Kopf Instruments). An injection needle (25 or 31 gauge) was vertically inserted into the MFC (+3.0 mm to Bregma, 1.0 mm to midline, -1.5 mm from pia mater). Muscimol solution was injected via the microsyringe pump at 0.2 μl/min by totally 0.5 ul in each hemisphere. After injections were completed, the needle was left for 5 minutes to allow for diffusion and then slowly retracted. Then, a silicon probe was inserted into DS of left hemisphere as described above. Then, the rats were tested the behavioral task 1 hour after end of muscimol infusion. For control group, multi-neuron and juxtacellular recordings were conducted.

### Histology

Animals were deeply anaesthetized with Urethane (2-3 g/kg, i.p.) and then perfused intracardially with 0.9% chilled saline followed by 4% paraformaldehyde (PFA) dissolved in 0.1 M phosphate buffer (PB). The fixed brain was stored in 4% PFA overnight and then stored in 30% sucrose solution in 0.1 M PB over 2 weeks. Postfixed brains were frozen and coronally sliced into 50-μm thick serial sections with a microtome Cryostat (HM500OM, Microm). The brain sections were stored in 0.1 M PB at 4°C overnight. The brain sections were subject for immunostaining to detect FG and Neurobiotin (Nb). For fluorescent visualizations of FG labelled neurons and Nb-loaded neuron, the brain sections were incubated with a rabbit antibody of FG (AB153, 1:3000 dilution, Millipore) at 4°C overnight, followed by incubation with a goat anti-rabbit antibody of FG conjugated with Alexa-594 (A11012, 1:500 dilution, Invitrogen) together with antibody Strept Avidine Alexa-488 (A11012, 1:250 dilution, Invitrogen) of Nb over 2 hours. The slices were washed with PB and mount on a slide glass and dried in a shaded box overnight. Fluorescent images were imaged with a fluorescence microscopy (Olympus, AX70) in order to check if FG labeled neurons were observed around the silicon probe recording locations in MFC and near the FG injection site in DS as well as if the Nb-loaded neurons were observed in MFC. After the fluorescent imaging, the brain sections were re-stained the Nb-loaded neuron using the avidin-biotin-horseradish peroxidase complex (Vectastain Elite ABC; Vector Laboratories, 1:200 dilution) with diaminobenzidine and nickel as demonstrated previously [23,54]. Finally, the slices were counterstained with Neural Red Nissl, and observed with a microscopy to check the tracks (scars) due to probe insertion. We judged the recording locations in MFC and DS and the AP coordinate of brain sections according to the rat brain atlas [56].

### Data analysis

All of behavioral and neuronal data were analyzed by custom-written MATLAB scripts (The MathWorks, Inc.). We analyzed behavioral data on 20 rats performing the task at high reward acquisition probability (>0.75) through 30 recording sessions.

#### Logistic regression analysis

We estimated effects of past outcome and past choice on current choice (repeat or switch) by the following regression model [14].

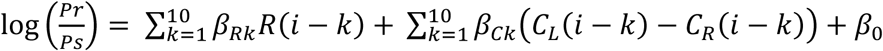

where *p*_r_ (or *p*_s_) is the probability of repeat choice (or switch choice) in the *i*th trial. The variables R(*i*) and C_L_(*i*) (or C_R_(*i*)) indicate presence of reward delivery (0 or 1) and left (or right) choice (0 or 1) in the *i*th trial, respectively. The coefficient β_R*k*_ and β_C*k*_ represent the effect of past rewards and choices, respectively. β_0_ is an intercept. These coefficients were calculated at each session. We checked if the coefficient was significantly different from zero by t-test with Bonferroni correction (Fig. 1d).

#### Spike sorting, clustering, and refining

Spike event from multiunit (or juxtacellular) activity was isolated by a custom-made semi-automatic spike-sorting program EToS [12 (or 5) feature dimensions for 4 (or 1) channels; high-pass filter, 300 Hz; time-resolution, 20 kHz; spike-detection interval, >0.5 ms) [57,58]. The sorted spike clusters were combined, divided, and discarded manually to refine single-neuron clusters by Klusters [59]. To avoid overlapping of detection of same units recorded among spatially distinct tetrodes, we checked cross-correlations of spike times among isolated units across all of tetrodes. If there was a high correlation peak only at zero time between a pair of units, one of the units was excluded from further analyses because those spikes which originated from the same neuron were presumably recorded through different tetrodes.

For comparison of neural activity between MFC and DS, we analyzed behavioral and electrophysiological data (isolated cells from multi-neuron activity) in 12 sessions from 10 rats: experiment-id (reward acquisition probability, the number of well isolated cells), R982-r1 (p = 0.828, MFC/DS = 45/43), R983-r1 (p = 0.837, MFC/DS = 35/55), R983-r2 (p = 0.834, MFC/DS = 58/48), R985-r1 (p = 0.853, MFC/DS = 30/60), R985-r2 (p = 0.867, MFC/DS = 66/26), R986-r1 (p = 0.813, MFC/DS = 16/20), R991-r1 (p = 0.806, MFC/DS = 46/31), R1000-r1 (p = 0.784, MFC/DS = 12/32), R1004-r1 (p = 0.792, MFC/DS = 20/51), R1005-r1 (p = 0.794, MFC/DS = 65/64), R1009-r1 (p = 0.774, MFC/DS = 35/33), R1012-r1 (p = 0.755, MFC/DS = 40/26). To examine effects of muscimol injection in MFC on neuronal activation of DS cells, we analyzed data from 130 and 77 DS cells of the 3 rats in muscimol group at the first and second recording sessions, respectively.

#### Event-related neuronal activity

Task-event-related neuronal activity was examined on well isolated units. To make peri-event time histograms (PETHs), we calculated mean and SEM of instantaneous firing rate (FR) in a 20-ms bin around task events (Start cue onset, Go tone onset, Choice response, and Reward delivery) and the PETHs were smoothed with a Gauss filter (SD = 40 ms). To compare firing pattern across neurons and recording sessions, the PETHs were transformed the FR into z-score by using the mean and SD of control activity, which were FRs in 200-ms at pseudorandom times (1000 time points) in each recording session. Then, the z-scores were normalized by the absolute peak value (Fig. 3e).

#### Neuronal selectivity for choice and outcome

Statistical significance of neuronal modulation by choice and outcome was analyzed by a two-way ANOVA with choice (left and right) and outcome (reward and no-reward) factors (p <0.05) for each FR in 0.2-s wide sliding window by 20 ms increment around choice response (±3 s) (Fig. 3f). To quantify the preference of selectivity for choice and outcome, we used a receiver-operating characteristic (ROC) analysis for neurons showing the statistical significance by two-way ANOVA at each window [60]. The proportion of trials for a condition (left choice or rewarded trials) was plotted against that of trials for another condition, (right choice or unrewarded) while changing a criterion for the FR. If the area under the plotted ROC curve (AROC) was 0.5 (0 or 1), the two conditions are indistinguishable (or completely different, respectively). If AROC was larger (smaller) than 0.5, FR is larger in right choice (left choice) or unrewarded outcome (rewarded outcome) than under the opposite condition. To compare the selectivity for choice and outcome modulated activity between MFC and DS, the AROC was transformed as a selectivity index. Selectivity index = |AROC – 0.5| + 0.5 (Fig. 3g).

#### Analysis of neural trajectory separation

All analyses for population activity were separately performed on data sampled at individual recording session. Fisher’s linear discriminant [35] was used to find the degree of discrimination between neural trajectories classified into two conditions (e.g., Left and Right choices). To define a Ch-hyperplane, we constructed the distributions of the corresponding ***r***(*t*), which is *N*-dimensional population FR vector (N corresponds to the number of population of neurons), separately for Left and Right choice trials in each 0.2-s sliding window by 50 ms increment time around choice response (±3 s). Our task is to find a hyperplane, or equivalently the normal vector **w** of this hyperplane, that best divides the two distributions projected onto the direction of **w**. If the two distributions have the means and covariance matrices **m**_1_, Σ_1_ and **m**_2_, **Σ**_2_, respectively, we can obtain **w** by maximizing the ratio of the between-class variance to the within-class variance,

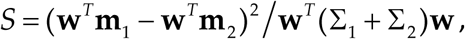

where **w**^T^**m**_1_ and **w**^T^**m**_2_ are the means of the two projected distributions and **w**^T^**Σ**_1_**w** and **w**^T^**Σ**_2_**w** are their variances. We can show that *S* is maximized if **w** ∝(Σ_1_ + Σ_2_)^−1^(**m**_2_ − **m**_1_). The coefficient of proportion can be determined by the normalization condition: | **w** |= 1.

Thus, we can calculate the discrimination function between Left and Right choice trials as 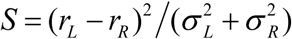 from the means (*r*_*L*_, *r*_*R*_) and standard deviations (*σ*_*L*_, *σ*_*R*_) of the trial-by-trial firing rates in these trials. The discrimination degree between the two clusters is defined as *d’* = *S*.

The ***r***(*t*) in each 0.2-s wide window (0.2 s increment time) around choice response (±3 s) was projected onto the axis orthogonal to the hyperplane by calculating inner product of ***r***(t) and the normal vector **w**_choice_, which yielded maximal degree of discrimination (*d’*). For instance, population rate at time *t* vectors was projected onto the axis orthogonal to “Ch-hyperplane”, *v*_*t*_ = < ***r***(*t*), ***w***_choice_>. In this case, we termed the projected rate vector “Ch-projected trajectory”. We performed the same procedure to find “Ot-hyperplane” (**w**_outcome_) by classifying trials into rewarded and unrewarded conditions, and to obtain the projected rate vector “Ot-projected trajectory”.

#### Mutual information

To quantify the difference in Ch-projected trajectory (or Ot-projected trajectory), we calculated the mutual information [61] between choices (or outcomes) and the projected population rate vectors *v*_*t*_:

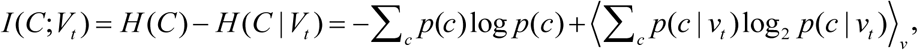

where *C* and *V*_*t*_ denote the set of choices (or outcome) *c* and the set of projected population activity *v*_*t*_, respectively, and *p*(*c*) is the choice probability (or outcome probability), *p*(*c*|*v*_*t*_) is the conditional probability of choice (or outcome) *c* when *v*_*t*_ is observed at time *t*, and the parenthesis means an averaging over the values of *v*_*t*_. The mutual information was calculated at each time window (bin = 0.2 s) around choice response (±3 s), and was normalized by the total entropy of choice *H*(*C*) (or total entropy of outcome) to provide the percentage of information as demonstrated before [12].

#### Pairwise correlation analysis of population activity

A Pearson correlation was performed for Ch-projected trajectory and Ot-projected trajectory between MFC and DS at each time window (bin = 0.2 s) around choice response (±3 s). Because the correlations are sensitive to the number of trials, we randomly selected 431 trials, which was the minimum total number of trials among 9 sessions, and calculated correlation coefficient so that the correlation coefficient was calculated with the same total number of trials sampled. We repeated this procedure 10 times to obtain mean correlation coefficients (Fig. 4e and Fig. S4). To compare the correlations of trajectories between rewarded and unrewarded trials, we calculated correlation coefficient by randomly selected 63 trials, which was the minimum total number of unrewarded trials among 9 sessions, for each outcome condition (only rewarded trials, only unrewarded trials, trials without distinction). We repeated this procedure 10 times to obtain mean correlation coefficients (Fig. S5b).

#### Cross-correlation analysis of spike times

We calculated cross-correlograms (CCGs) under 3 conditions (left choice trials, right choice trials, and rewarded trials) for all MFC-DS cell-pairs. We used spike times in a 5-s wide window centered on a specific time at which a maximum absolute correlation of neural trajectories was yielded at Ch-hyperplane (or Ot-hyperplane) referred as window Ch (or window Ot) (Fig. 4e and Fig. S4). As a control, we calculated task-event irrelevant CCGs by randomly sampling X time points within the same recording session for the same cell-pair. X corresponds to the number of trials used for calculation of task-event relevant CCGs under each condition. For example, the CCG between MFC and DS cells was calculated by counting spikes of MFC cell in a window ranging from -0.1 to +0.1 s relative to each spike time of DS cell (bin = 1 ms). Spike synchrony was detected by comparing the peak value in CCGs within ±20 ms with the mean+4SD of values in a baseline window (−100 to -80 ms and +80 to +100 ms) under each condition. If the peak value was larger than this criterion, we judged that the neuron pair showed spike synchrony. We did not assess the spike synchrony if the mean of baseline values was less than 1 or the SD was 0 due to scarce spiking. We did not look into negative spike correlation. For each cell-pair showing significant CCG peak values, we also calculated CCGs by shuffling trials between MFC and DS cells (Fig. S6). To compare peak value of CCGs, each CCG was z-scored by the mean and SD of the baseline values (Fig. S6 and 5b).

