## Supplemental text for "Concomitant processing of choice and outcome in frontal corticostriatal ensembles correlates with performance of rats"

### *Change in firing rates along with repetitive choice or outcome history in single neurons in MFC and DS*

A small number of MFC and DS neurons revealed dynamic change in firing rates (FR) along with repetitive choices or outcome history (Fig. S2b). Statistical significance of neuronal modulation by choice position and repetitive choices was judged by a two-way ANOVA with choice position (left and right) and repetitive choice (the number of repeating same choice after rewarded trial or unrewarded outcome) factors ($p < 0.05$) for each FR in 0.2-s wide sliding window by 20 ms increment around choice response (±3 s). For instance, choice-modulated activation dynamically changed as rewarded choice repeated or depending on action selection (i.e., repeat or switch) even if the choice position was identical (Fig. S2b, left). The MFC neuron revealed clear choice-position selective activation, but the discharged rate was modulated by repetitive choice pattern. A DS neuron fired more after response until reward delivery but also during the late epoch of outcome period in unrewarded trials (Fig. S2b, middle). Another MFC neuron revealed sparse firing during the trial in switching choice, but afterwards increased discharge rate after repeating rewarded left choice trials (Fig. S2b, right).

***Degree of discrimination in population activity***

We examined how neural populations in MFC and DS encode information about decision making in each session. We explored the time at which the neural trajectories encoding left and right choice decisions were maximally separated by using Fisher's linear discriminant. Discriminability index *d'* describes the distance between categorized rate vectors measured orthogonally to such a hyperplane. The index *d'* for outcome (Ot) hyperplane in both MFC and DS reached a maximum distance after variable reward delivery time (2nd Delay), and this temporal pattern was consistently observed in all sessions. On the other hand, temporal pattern of *d'* for choice (Ch) hyperplane was not consistent among sessions and between MFC and DS (Fig. S3a). The peak values of *d'* for Ch- and Ot-hyperplanes were correlated to the dimensionality (i.e. *N*) of population vector (Pearson correlation analysis, Ch-hyperplane, MFC, $r = 0.739$, $p = 0.0060$, DS, $r = 0.630$, $p = 0.0280$; Ot-hyperplane, MFC, $r = 0.616$, $p = 0.0327$, DS, $r = 0.612$, $p = 0.0344$) (Fig. S3b). In particular, the peak of d' for Ch-hyperplane was not clear when number of MFC neurons was smaller than 25. Therefore, to keep a sufficient number of cells for evaluating *d'*, we used only 9 dataset (R982-r1, R983-r1, R983-r2, R985-r1, R985-r2, R991-r1, R1005-r1, R1009-r1, R1012-r1), which includes the large number of neurons ($n \geqq 25$) in both regions per session, in the present analyses.
